# Transcriptomic prey-capture responses in convergently evolved carnivorous pitcher plants

**DOI:** 10.1101/2025.07.16.665073

**Authors:** Takanori Wakatake, Kenji Fukushima

## Abstract

The Australian pitcher plant *Cephalotus* and the Asian pitcher plant *Nepenthes* exhibit striking morphological and functional similarities, serving as compelling examples of convergent evolution. Although trapping pitchers in both lineages represent some of the most elaborate leaf structures in angiosperms, it remains unknown whether their analogous phenotypes share common molecular foundations, especially at the level of gene expression. Here, we conducted tissue-specific RNA-seq experiments coupled with feeding treatments in *C. follicularis*, mirroring the available expression dataset of *N. gracilis* to analyze gene expression evolution underlying the phenotypic convergence. Functionally equivalent tissues in the two species tended to express similar gene sets, with common transcriptional responses that activate amino acid metabolism and protein synthesis upon the feeding treatment, yet with distinct transcriptional regulation of digestive enzyme genes. Additionally, we found multiple cases of combined convergence in expression and protein sequences in genes preferentially expressed in gland-containing tissues. Our study showcases how common and unique transcriptional components are integrated to shape the independent emergence of complex leaf structures in angiosperms.

## Introduction

Convergent evolution, wherein distinct lineages independently evolve similar solutions to ecological pressures, offers insights into how organisms respond to natural selection. Among the most captivating examples are carnivorous plants, which collectively evolved in six plant orders of angiosperms (Freund et al. 2022). These remarkable organisms have evolved mechanisms for nutrient acquisition through the capture and digestion of prey, a strategy distinct from the soil-based nutrient absorption typical of most land plants.

Most carnivorous plants have modified their leaves into trapping organs. To integrate trapping functions into leaves originally optimized for photosynthesis, considerable amounts of organ complexity have been added during carnivorous plant evolution, culminating in pitcher leaves, which represent one of the most intricate leaf structures observed in angiosperms (Tsukaya 2014). Small arthropods that slip into this cavity become ensnared and are degraded by the cocktail of digestive enzymes secreted from glands. The released nutrients are absorbed from glands and then distributed throughout the plant body for growth and reproduction.

Caryophyllales encompasses a diverse and ancient lineage of carnivorous plants, yet pitfall traps evolved exclusively within *Nepenthes*. *Nepenthes* develops trapping pitchers in the distal half of a single leaf, which bears a tendrilled leaf-blade-like structure in the basal half. Thus, a single *Nepenthes* leaf is segmented along the proximodistal axis, featuring distinctive tissues each specialized for photosynthesis, carnivory, and its sub-functions such as digestion. Gland cells on the inner wall of the digestive zone at the pitcher’s base serve the dual purpose of secreting digestive enzymes and absorbing nutrients released during prey digestion (Freund et al. 2022). Above the digestive zone, the waxy zone hinders trapped prey from escaping, thanks to a slippery crystallized epicuticular wax (Gorb et al. 2005). The lid and the pitcher’s rim, the peristome, secrete nectar to attract prey (Moran 1996). The peristome also plays an important role in capturing prey by providing a slippery foothold for insects with wettable microscopic and macroscopic surface structures (Bohn and Federle 2004; Labonte et al. 2021). Nutrients absorbed within the pitcher leaf are transported to the entire plant body through the tendril, which connects the pitcher to the leaf-blade-like tissue (flat part).

Despite evolving independently in distant lineages, *Cephalotus follicularis*, a carnivorous plant in the Oxalidales order and the exclusive member of the Cephalotaceae family, shares a remarkably similar structural and functional configuration of trapping pitchers with *Nepenthes* (Fig. 1a,b). *Cephalotus* develops carnivorous pitcher leaves alongside photosynthetic flat leaves depending on ambient temperature (Fukushima et al. 2017). The lower part of the pitcher leaves houses two distinct types of secretory glands (Freund et al. 2022). The small glands are densely clustered in glandular patches, which are situated symmetrically near the bottom of the pitcher leaves and can be distinguished by their colors and tissue thickness. The glandular patches are surrounded by a higher density of the large glands that are also distributed to the upper inner wall with gradually reduced density (Fig. 1b) (Juniper et al. 1989). The downward-pointing hairs on the neck prevent prey from escaping the cavity (Juniper et al. 1989). A roof-like lid features nectary glands on its bottom side (Juniper et al. 1989; Vogel 1998). With a few distinctions, such as gland morphology and distribution, *Cephalotus* pitchers share many hallmark characteristics with *Nepenthes* pitchers.

**Fig. 1.**
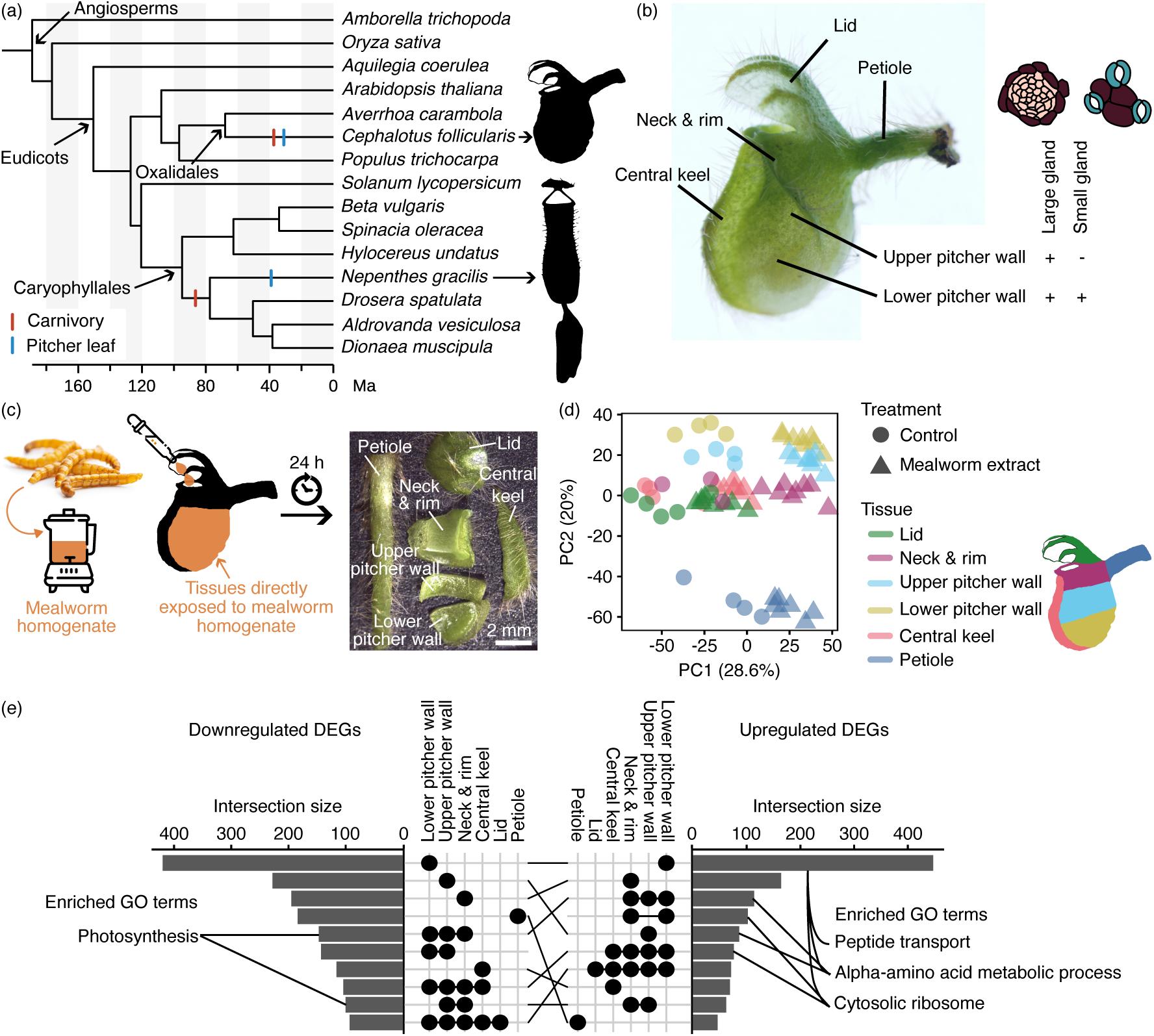
*Cephalotus* pitcher leaves respond to the feeding treatment. (a) The phylogenetic tree including 15 angiosperms used for analysis in this study. Color bars indicate the independent origins of plant carnivory and pitcher leaves in the Oxalidales and the Caryophyllales. The placement of the bars on the phylogenetic tree does not correspond to the estimated evolutionary timing of the trait’s origin. (b) The tissue structure of the *Cephalotus* pitcher leaves. The positions of the two types of glands were indicated. (c) Illustration of the RNA-seq experiment coupled with the mealworm extract treatment. (d) Principal component analysis of expression values of 3,987 differentially expressed genes (DEGs) between control and worm extract-treated samples. Points correspond to RNA-seq samples. (e) The upset plot shows the number of up-regulated (right panel) and down-regulated (left panel) genes in each set association in the middle. Selected GO terms enriched in the set association are indicated. The same set associations in the two upset plots were connected with black lines. Image credits: the mealworm photo and icons from freepik.com. The photographs of the *Cephalotus* pitcher in (b) and (c) were reproduced from the previous publications (Fukushima et al., 2021; Saul et al., 2023).

Carnivorous plants often exhibit a feeding response, which expedites prey digestion and nutrient absorption following prey capture. This response typically involves the reduction of pH in digestive fluids, enhancing the proteolytic activity of these fluids, the secretion of digestive enzymes, and the activation of nutrient absorption. Various cues can trigger this feeding response in distinct ways. For instance, in *Dionaea muscipula* (Venus flytrap) having snap trap, touch-induced action potentials, repeated mechanical stimuli from struggling trapped prey, and chemicals released from degraded prey trigger trap movement, digestive enzyme secretions, and nutrient absorption (Libiaková et al. 2014; Bemm et al. 2016; Böhm et al. 2016; Pavlovič et al. 2017). In contrast to *Dionaea*, pitcher leaves do not rely on mechanical stimuli and use chemical cues to detect the presence of prey (Gallie and Chang 1997). Chitin, a primary component of insect exoskeletons, as well as ammonium released during protein digestion and proteins such as bovine serum albumin (BSA), each induce distinctive feeding responses in *Nepenthes* (Yilamujiang et al. 2016; Saganová et al. 2018). BSA and ammonium strongly induce digestive activities also in *Sarracenia purpurea*, a carnivorous pitcher plant in Ericales (Gallie and Chang, 1997).

In this study, we generated a tissue-specific RNA-seq dataset for *Cephalotus* under feeding treatment conditions and paired it with a mirrored dataset from *Nepenthes* (Saul et al. 2023). This approach enabled a direct comparison of transcriptomic features between these two deeply diverged, independently evolved pitcher plant lineages. Our analyses uncovered shared expression profiles as well as contrasting regulatory responses associated with prey capture. In addition, we identified convergent amino acid substitutions in 11 proteins upregulated or specifically expressed in gland-containing tissues, suggesting potential adaptive evolution at both expression and protein sequence levels. We discuss the evolutionary implications of these shared and divergent molecular features that underpin the complex pitcher organizations.

## Results and Discussion

### Prey-induced transcriptional responses of *Cephalotus* pitcher tissues

To explore how *Cephalotus* responds to prey capture at the transcriptome level, we performed an RNA-seq experiment with a feeding treatment. To mimic prey capture, we supplied pitchers with the water extraction of dried mealworms. As a negative control, water was supplied. After 24 hours, the treated pitchers were dissected into six parts (lid, neck and rim, upper pitcher wall, lower pitcher wall, central keel, and petiole) to examine tissue-specific responses, following the dissection scheme established in the previous publication (Saul et al. 2023), and the dissected tissues were subjected to the RNA-seq experiment (Fig. 1c). To overview the dataset, a principal component analysis (PCA) was performed using differentially expressed genes (DEGs) that responded to the feeding treatment in at least one tissue (3,798 genes, false discovery rate (FDR) < 0.05, |log_2_ fold change| ≥ 1.0) (supplementary table S1). The first principal component (PC1) explained 28.6% of the variation, clearly separating control and fed samples in all tissues (Fig. 1d). The pitcher tissues were separated mainly with the PC2 axis. The number of unique DEGs (i.e., genes differentially expressed only in the focal tissue) was highest in the lower pitcher wall (Fig. 1e), where two types of glands are located (Fig. 1b). In this tissue, GO terms ‘peptide transport’, ‘alpha-amino acid metabolic process’, and ‘cytosol ribosome’ were significantly enriched in the up-regulated DEGs, suggesting a role in absorbing and assimilating proteinaceous degradation product and protein synthesis (Fig. 1e) (supplementary table S2). The term ‘alpha-amino acid metabolic process’ was also enriched in the upregulated DEGs in the upper pitcher wall, in correlation with the location of the large glands. ‘Cytosol ribosome’ showed enrichment in all analyzed tissues but the petiole (Fig. 1e), suggesting that protein synthesis is activated in the whole pitcher in response to the feeding treatment but is decoupled in the petiole. Among the commonly downregulated genes, we found that the GO term ‘photosynthesis’ was enriched in the pitcher-forming tissues (Fig. 1e). While prey capture generally enhances photosynthesis in carnivorous plants over the long term (2 weeks to 5 months) by increasing nutrient availability (Givnish et al. 1984; Farnsworth and Ellison 2008; Pavlovič et al. 2009; Pavlovič et al. 2011; He and Zain 2012), our results indicate a more immediate transcriptional response that prioritizes carnivory over photosynthesis. This response appears mechanistically distinct from that observed in *Di. muscipula*, where photosynthetic rates rapidly decline upon trigger hair stimulation but recover within 10 minutes (Pavlovič et al. 2010). The gene expression changes in *Cephalotus* suggest a more sustained downregulation, potentially reflecting a resource allocation strategy that enhances digestive processes.

### Feeding responses in the two pitcher plant lineages

We previously conducted tissue-specific RNA-seq experiments with the same feeding treatment in the pitcher leaves of *N. gracilis*, a carnivorous pitcher plant that evolved separately from *Cephalotus* (Fig. 1a) (Saul et al. 2023), providing an opportunity to test whether independently evolved carnivorous pitcher leaves exhibit similar transcriptome responses upon prey capture. To enable direct comparison, we reanalyzed the *N. gracilis* dataset using the same method as for *Cephalotus*, and used the resulting output for downstream analyses (supplementary table S3). In contrast to *N. gracilis* pitcher, in which DEGs were detected almost exclusively in the digestive zone, *Cephalotus* showed DEGs in all analyzed pitcher tissues (Fig. 2a). The number of DEGs was the highest in the lower pitcher wall, which, together with the upper pitcher wall, was directly exposed to the mealworm extract. DEG abundance decreased progressively with the increasing distance from the treated site and reached its lowest in the most distant tissues, the petiole and the lid, though it never dropped entirely to zero (Fig. 2a). To integrate the gene expression between the two pitcher plant species split more than 100 million years ago (Saul et al., 2023), gene orthology was inferred using OrthoFinder with the inferred species tree (Fig. 1a) to inform phylogenetic relationships (supplementary table S4) (Emms and Kelly 2019). We defined the orthogroup-level feeding response as either up, down, or conflict by majority decision of member DEGs within orthogroups for each pitcher tissue (Fig. 2b) to evaluate the overlapped responses between the digestive zone of *N. gracilis* and all pitcher tissues of *C. follicularis* (Fig. 2c). The lower pitcher wall showed the largest number of orthogroups consistently upregulated in the digestive zone in *N. gracilis*, exhibiting the highest Jaccard similarity coefficient, which measures the degree of overlap between two orthogroup sets (Fig. 2d,e). The large portion of enriched GO terms in these shared orthogroups was related to ribosome (supplementary table S5). These analyses suggested a shared response to the feeding treatment in two phylogenetically distant pitcher plants.

**Fig. 2.**
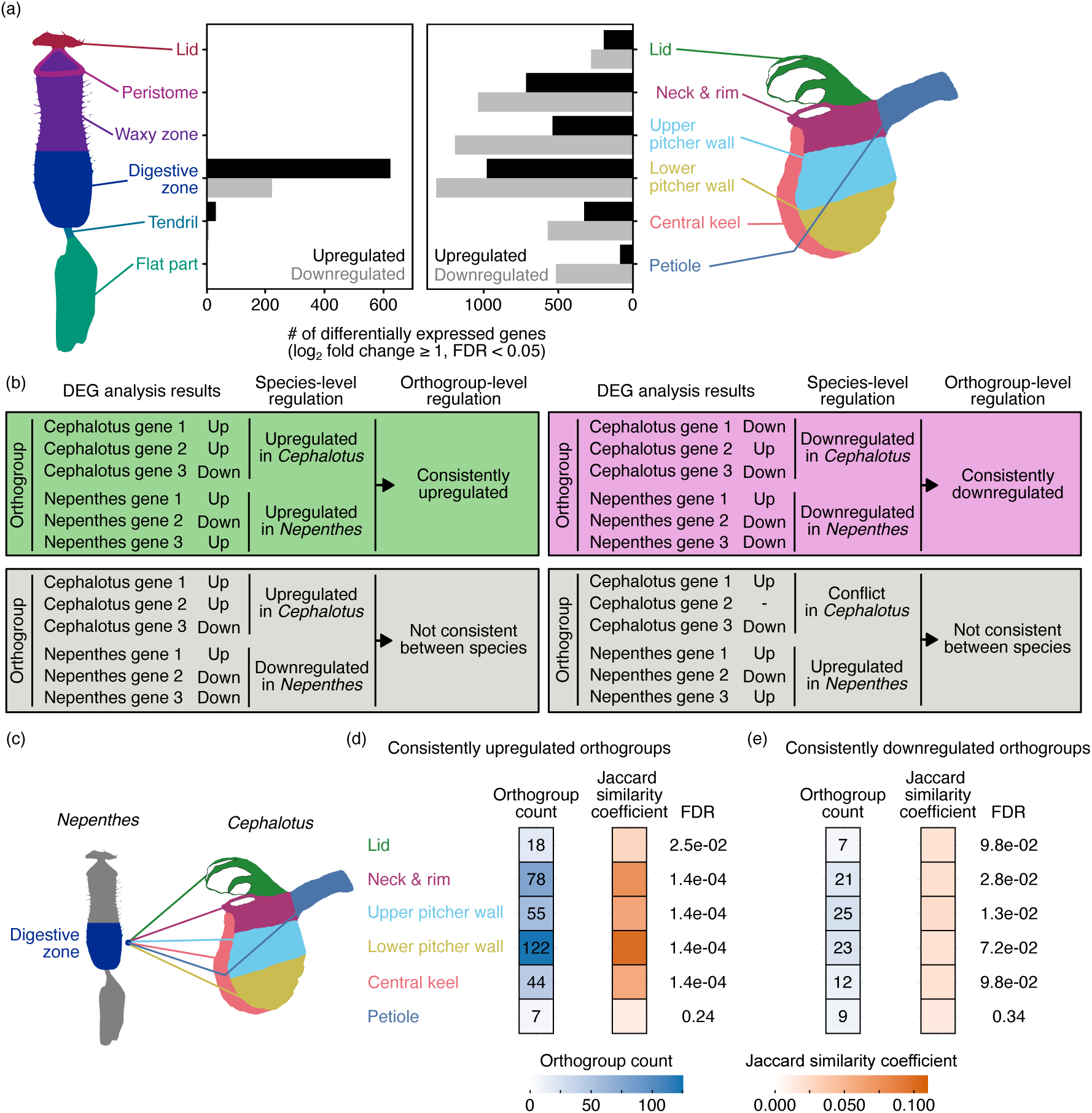
*Cephalotus* pitchers upregulate the gene set similar to *Nepenthes* pitchers in response to the feeding treatment. (a) The number of differentially expressed genes (DEGs) in each tissue in *N. gracilis* (left) and *C. follicularis* (right). Different tissues are indicated with different colors in the illustration. (b) The definition of the consistently upregulated (green) and downregulated (magenta) orthogroups in two species. Two examples with inconsistent regulation (gray) are shown in the second row. (c) Schematic comparison of DEGs between the digestive zone of *N. gracilis* and all pitcher tissues of *C. follicularis*. (d, e) Consistently upregulated and downregulated orthogroups were analyzed, respectively. The heatmaps indicating the number of shared orthogroups and the Jaccard similarity coefficient (JC, middle), and false discovery rate (FDR) values for JC (right) were shown. The numbers in the tiles indicate the shared orthogroup count between two groups. FDR values were calculated using the Benjamini-Hochberg method for multiple testing correction, following *P* value computation via permutation tests.

### Tissue-specific gene expression in the two pitcher plant lineages

Distinct tissues contribute to the various subfunctions that underpin the highly organized pitcher leaves. To compare tissue-specific basal gene expression levels in the two pitcher plant lineages, we applied self-organizing map (SOM) clustering to RNA-seq datasets under the control conditions. Genes with a high coefficient of variation in gene expression (top 10%, 1,750 genes for *C. follicularis*, 2303 genes for *N. gracilis*) were subjected to SOM clustering in a 4 × 3 grid with hexagonal topology (supplementary table S6).

In *N. gracilis*, cluster 4 exhibited high and specific expression in the digestive zone, and included nine out of the 18 genes orthologous to those encoding digestive fluid proteins in congeneric species (*N. alata, N. rafflesiana, and N. mirabilis*) (Fig. 3) (Saul et al., 2023). Similarly, in *C. follicularis*, cluster 5 exhibited high expression specifically in the upper and lower pitcher walls, and included 11 out of 12 genes encoding previously identified digestive fluid proteins (Fig. 3). Both clusters were enriched in GO terms associated with digestive functions, such as ‘chitinase activity,’ ‘aspartic-type endopeptidase activity,’ and ‘acid phosphatase activity,’ suggesting a functional link to the digestive processes (supplementary table S7). Cluster 5 of *C. follicularis* additionally exhibited enrichment for defense-related GO terms (e.g., ‘defense response to insect’ and ‘defense response to fungi’), likely reflecting an overlap between digestive physiology and immune functions.

**Fig. 3.**
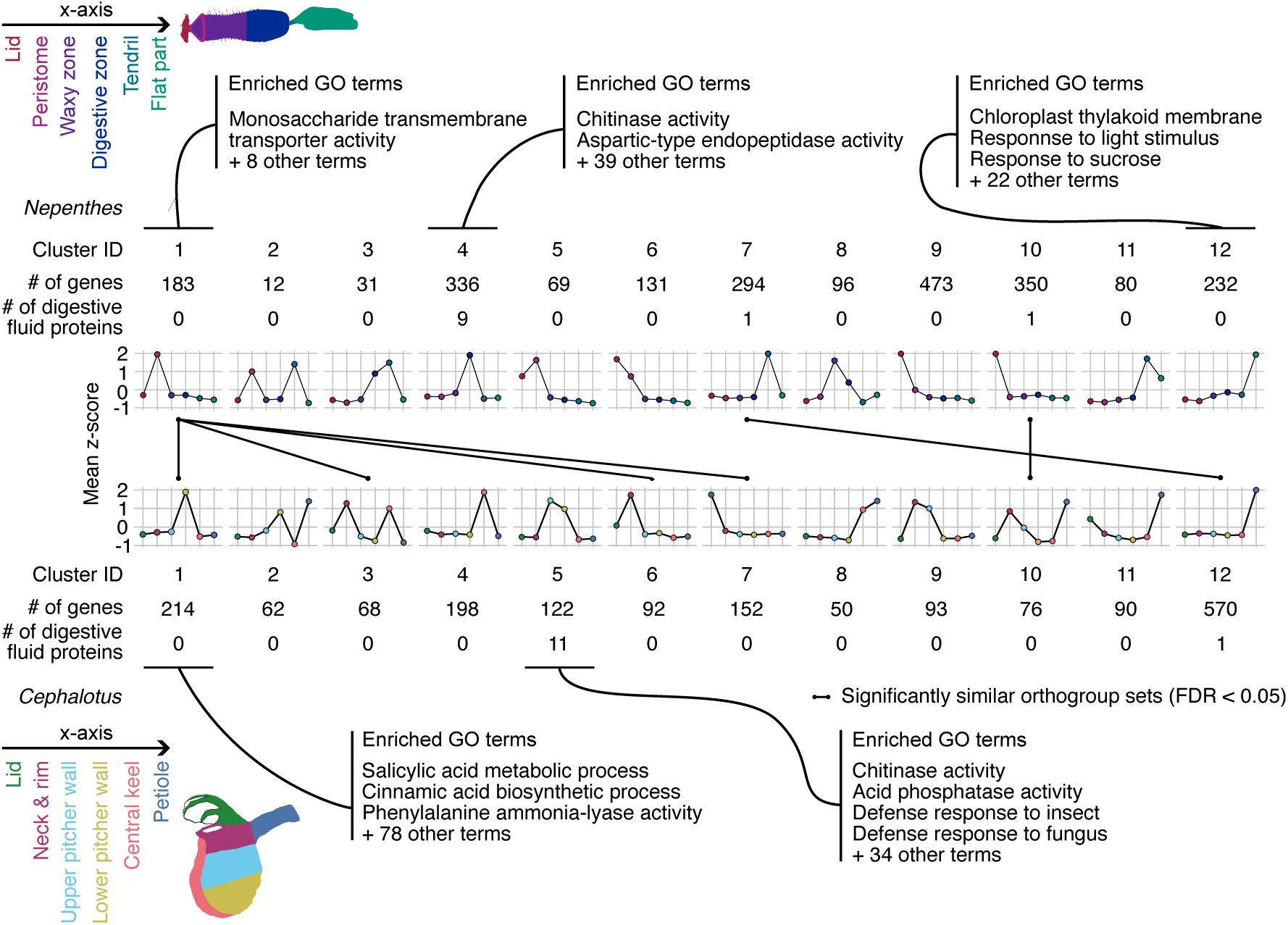
Comparison of tissue-specific gene expression under the basal conditions in *N. gracilis* and *C. follicularis* pitchers. SOM clustering was performed on genes with high coefficient of variation in expression across six pitcher tissues in *N. gracilis* (top) and *C. follicularis* (bottom). The x-axis labels are shown at the top-left and bottom-left corners of the panels. The y-axis represents the average z-score of gene expression within each cluster. Clusters with significantly similar orthogroup compositions between the two species (Jaccard similarity coefficient, FDR < 0.05) are connected by black lines. Selected enriched GO terms for representative clusters are also shown. FDR values were calculated using the Benjamini–Hochberg method for multiple testing correction after *P* values were obtained through permutation tests.

Distinct clusters were also associated with other specialized pitcher tissues. In *N. gracilis*, cluster 1, which was predominantly expressed in the peristome, was enriched in GO terms related to ‘monosaccharide transmembrane transporter activity,’ consistent with a role in nectar secretion. In addition, cluster 12, which was highly expressed in the basal flat part of the *Nepenthes* leaves, was enriched in GO terms related to photosynthesis and light response, consistent with its photosynthetic role. In contrast, *C. follicularis* cluster 1, which showed specific expression in the lower pitcher wall including the glandular patch, was enriched in GO terms related to plant hormone metabolism, such as ‘salicylic acid metabolic process,’ ‘auxin biosynthetic process,’ and ‘cytokinin metabolic process.’ While the enrichment of salicylic acid metabolism genes suggests a possible regulatory link between salicylic acid (SA) signaling and the expression of digestive fluid proteins, potentially mediated through immunity pathways (Bemm et al. 2016), auxin and cytokinin pathways may also have previously unrecognized links to carnivorous traits.

To quantitatively assess the similarity of tissue-specific gene expressions between the two species, we calculated the Jaccard similarity coefficient for orthogroup sets in a pair of SOM clusters from *N. gracilis* and *C. follicularis* (Fig. 3 and supplementary fig. S1). *Nepenthes* cluster 7, which showed high expression in the tendril, was most similar to *C. follicularis* cluster 12, which was highly expressed in the petiole. Both tissues connect the pitcher to the main plant body. Similarly, *N. gracilis* cluster 1, which showed peak expression in the peristome, displayed notable similarity to *C. follicularis* clusters 3 and 6, where expression peaked in the neck and rim tissues (Fig. 3). Unexpectedly, *N. gracilis* cluster 4, which exhibited high expression in the digestive zone, did not show significant similarity to *C. follicularis* cluster 5, despite their shared expression of genes encoding digestive fluid proteins. We extended the same SOM clustering analysis on the expression dataset after the feeding treatment (supplementary fig. S2 and supplementary tables S8,9), and calculated the Jaccard similarity coefficient (supplementary fig. S3). This revealed significant similarity between functionally equivalent tissues: the digestive zone of *N. gracilis* and the upper and lower pitcher walls of *C. follicularis*, and the flat part of *N. gracilis* and the central keel of *C. follicularis*. Together, these results indicate that functionally equivalent pitcher tissues tend to express similar gene sets. Furthermore, gene expression in certain tissues (e.g., the digestive zone and the upper and lower pitcher wall, and the flat part and the central keel) becomes more similar following feeding, suggesting a convergence in molecular responses to prey capture between the two species.

### Feeding responses of genes encoding digestive fluid proteins

One of the well-documented feeding responses in various carnivorous plants is the upregulation of secreted proteins in the digestive cocktail. We previously reported tissue-specific expression patterns of genes encoding digestive fluid proteins in *C. follicularis* under the control condition (Saul et al. 2023), however, their responses to prey capture remain unexplored. We examined the transcriptional responses of these genes and compared them with those in *N. gracilis*. In *C. follicularis*, all examined genes, except for *PRp27*, showed high tissue-specific basal expression in both the upper and lower pitcher walls, mirroring the distribution of the large gland (Fig. 4a). In *N. gracilis*, approximately half of the orthologs encoding digestive fluid proteins exhibited highly specific expression in the digestive zone even prior to feeding treatment (Fig. 4a). These expression patterns suggest constitutive protein production, consistent with the prevalence of digestive fluid proteins in unfed pitchers (Buch et al. 2015; Saganová et al. 2018; Dkhar et al. 2020). Basal expression levels of these genes were significantly higher in *C. follicularis* than in *N. gracilis* (Fig. 4b).

**Fig. 4.**
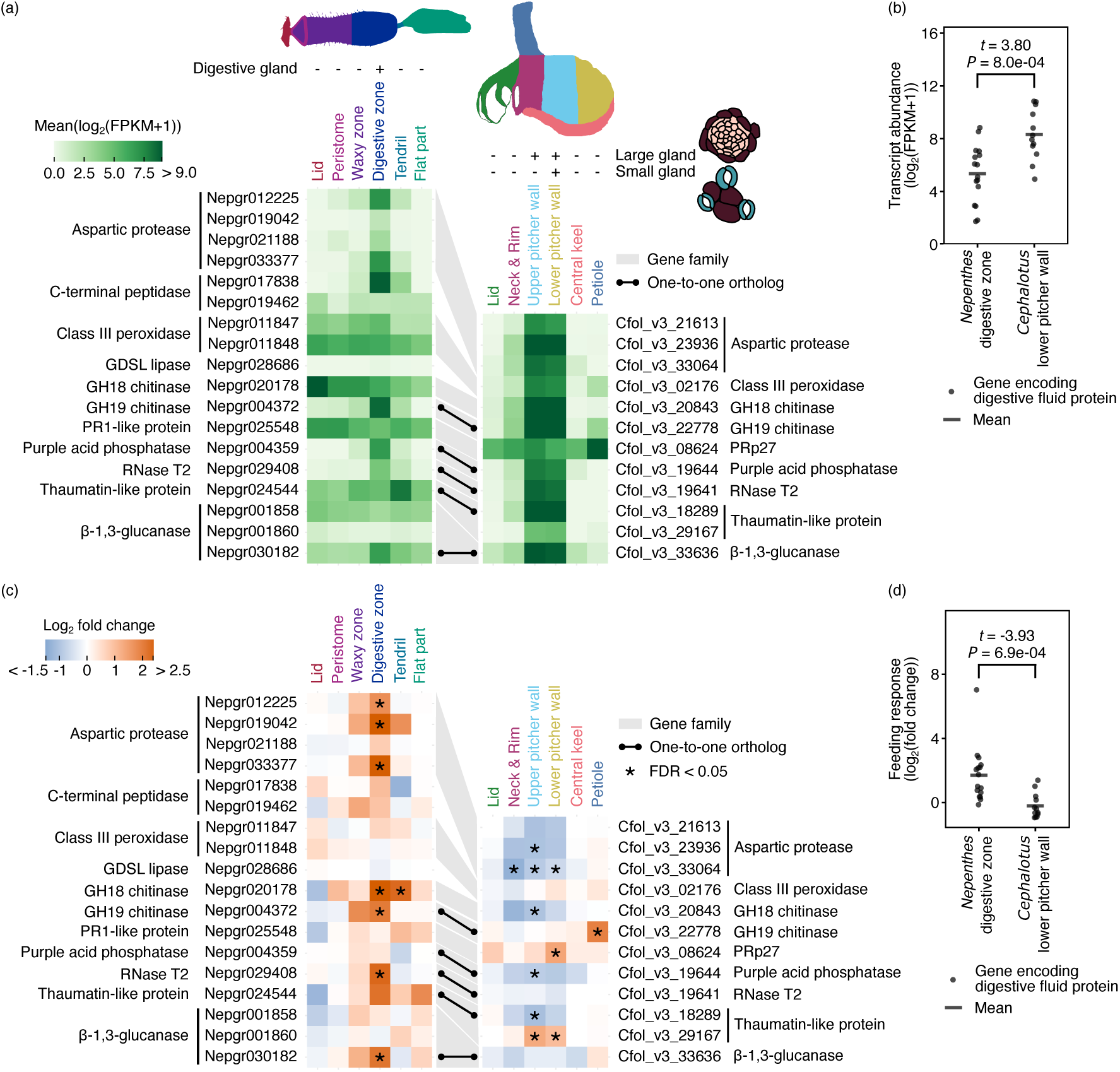
Differential response of genes encoding digestive fluid proteins to the feeding treatment in *Cephalotus* and *Nepenthes* pitchers. (a) Tissue-specific transcript abundance (log2(FPKM+1)) of genes encoding digestive fluid proteins. (b) Expression levels of the genes encoding digestive fluid proteins in the digestive zone and the lower pitcher wall before the feeding treatment. (c) Tissue-specific log2-fold change of genes encoding digestive fluid proteins upon the feeding treatment. (d) Log2-fold change of the genes encoding digestive fluid proteins in the digestive zone and the lower pitcher wall. Grey ribbons connect genes in the same gene family. The black dots and lines connect the orthologous genes. Asterisks in (c) indicate FDR < 0.05 in differential gene expression analysis. *P* values and *t* statistics of Welch’s two-sided *t*-tests are shown above the plot in (b) and (d).

Next, we compared the feeding responses of the genes encoding digestive fluid proteins. Low-expression genes (log_2_(FPKM+1) < 1.0) in gland-containing tissues both before and after the feeding treatment were excluded from these analyses. Surprisingly, in *C. follicularis*, these genes tended to be downregulated after the feeding treatment (Fig. 4c,d). In contrast, in *N. gracilis*, seven digestive fluid protein orthologs were upregulated specifically in the digestive zone following feeding treatment. These expression profiles underscore the distinct transcriptional regulations between the two pitcher plant lineages; In *N. gracilis*, genes encoding digestive fluid proteins are induced by feeding, whereas in *Cephalotus*, they exhibit constitutive expression before prey capture (Fig. 4b) and are downregulated after feeding (Fig. 4d).

Interestingly, in *C. follicularis*, genes encoding a precursor of immune-eliciting peptide and elicitor-producing apoplastic proteases showed expression profiles similar to genes encoding digestive fluid proteins, suggesting continuous production of immune elicitors (supplementary text 1). Also, while jasmonic acid (JA) signaling plays a central role in regulating digestive enzyme secretion in carnivorous Caryophyllales, our transcriptomic analysis suggests a more limited or divergent involvement of JA in *C. follicularis* (supplementary text 2). Together, these examples highlight lineage-specific regulatory mechanisms underlying carnivorous responses, despite their common reliance on defense-related enzymes.

The downregulation of the *Cephalotus* genes encoding digestive fluid proteins contrasts with the common strategy observed in many other carnivorous plants, where digestion is transcriptionally enhanced following prey capture. Notably, both proteolytic activity and aspartic protease abundance in the digestive fluid remain nearly constant for seven days post-feeding in *C. follicularis* (Pavlovič et al. 2024), indicating that transcriptional repression does not immediately impact the protein pool in the digestive fluid. The high basal expressions prior to feeding may buffer the digestive capacity in the early stages of prey breakdown. The observed downregulation may reflect a resource reallocation from digestion to nutrient assimilation following prey acquisition. Consistent with this idea, genes associated with nitrogen assimilation were upregulated after the feeding treatment (supplementary text 3). However, as our analysis was limited to a single time point, it remains unclear whether this downregulation is sustained or transient. Future time-course experiments will be necessary to clarify the temporal dynamics of digestive fluid protein expression.

### Convergent protein evolution in genes expressed in gland-containing tissues

Gene expression in specialized tissues, such as the gland-bearing structures of pitcher plants, may cause proteins to adapt to new cellular contexts, potentially leading to the coevolution of gene expression patterns and protein sequences. Indeed, *C. follicularis* and carnivorous Caryophyllales (including *Nepenthes*) exhibit convergent amino acid substitutions in digestive enzymes (RNase T2, GH19 chitinase, and purple acid phosphatase) that likely alter their biochemical properties for prey digestion (Fukushima et al. 2017; Fukushima and Pollock 2023). To investigate these combined molecular signatures of convergent evolution in pitcher plants, we searched convergent amino acid substitutions among genes expressed in gland-containing tissues in *C. follicularis* and *N. gracilis* using CSUBST (Fukushima and Pollock 2023). CSUBST calculates the *ω_C_* metric which estimates the rate of protein convergence. With the neutral expectation of 1.0, higher *ω_C_* values indicate accelerated rates of protein convergence. For this analysis, we selected 173 orthogroups containing genes upregulated in the digestive zone or in the lower pitcher wall (FDR < 0.05, log_2_ fold change > 1), or specifically expressed in tissues with glands (SOM cluster 4 for *N. gracilis* or SOM clusters 1 and 5 for *C. follicularis*) (supplementary dataset). With the thresholds for the number of convergent amino acid substitutions (*O_C_^N^* > 3.0) and for protein convergence rates (*ω_C_* > 3.0) to the branch pairs leading to *C. follicularis* and *N. gracilis* genes, we obtained 11 convergent branch pairs from 11 orthogroups (supplementary table S10). Although a different species set was used compared to the previous study (Fukushima and Pollock 2023), we successfully obtained high *ω_C_* values in RNase T2 (OG0001136) in *C. follicularis* and carnivorous Caryophyllales. In addition, the result included two proteins found in digestive fluids, S1-P1 nuclease and polygalacturonase-inhibiting protein (PGIP) (Schulze et al. 2012; Lee et al. 2016; Kocáb et al. 2020; Arai et al. 2021), and HIGH AFFINITY K^+^ TRANSPORTER 5 (HAK5) from the KT/HAK/KUP family, which functions in potassium uptake in the glands of *Di. muscipula* (Scherzer et al. 2015).

S1-P1 nucleases have been detected in the digestive fluids of multiple carnivorous plants across different lineages including *Di. muscipula*, *Drosera adelae*, *N. ventrata*, and *Pinguicula × Tina* (Schulze et al. 2012; Lee et al. 2016; Kocáb et al. 2020; Arai et al. 2021). All S1-P1 nucleases found in digestive fluids, except for one of two from *Dr. adelae*, are closely related to *Arabidopsis* ENDO2 (AtENDO2: AT1G68290). We detected accelerated protein convergence in ENDO2 orthologs in *Nepenthes* and *Cephalotus* (NgENDO2: Nepgr028074 and CfENDO2: Cfol_v3_09804). *NgENDO2* showed specific expression in the digestive zone and slightly upregulated after the feeding treatment (Fig. 5a). *CfENDO2* showed high tissue-specific expression in the upper and the lower pitcher walls similar to other genes encoding digestive fluid proteins (Fig. 5a). High *ω_C_* value was also obtained between *CfENDO2* and the stem branch of carnivorous Caryophyllales orthologs, suggesting that amino acid-changing substitutions in this gene are associated with two-step evolution, the emergence of carnivory and subsequent establishment of pitcher leaf organizations (Fig. 5a), as suggested for RNase T2 (Fukushima et al. 2017). *AtENDO2* is not induced by either JA or SA, but is induced by pathogen infection (Zhang et al. 2020), thus can be considered as a defense response gene. While its optimal pH is 6.0 and 7.0 for single-stranded RNA (ssRNA) and ssDNA cleavage, respectively (Ko et al. 2012), AtENDO2 shows relatively higher enzymatic activity at acidic pH compared to other genes within the same family in *A. thaliana* (Lesniewicz et al. 2013), suggesting suitability for the acidic environment of digestive fluid. A total of six convergent amino acid substitutions were located near the substrate binding pocket (Fig. 5a) (Chou et al. 2013). The K48R substitution is predicted to enhance catalytic efficiency toward ssDNA and double-stranded DNA (dsDNA), as a reverse substitution (corresponding to R74K in SmNuc1 [GenBank: WP_005410840.1]) at this position has been reported to reduce catalytic efficiency to approximately 80% and 70% for ssDNA and dsDNA, respectively (Adámková et al. 2024). The Y59F substitution is located at the nucleotide binding site and can probably alter the pH optimum (Koval and Dohnálek 2018). S1-P1 nuclease DAN1 (NCBI accession: LC699681.1), an AtENDO2 homolog in a carnivorous Caryophyllales species *Dr. adelae*, is expressed specifically in the glandular tentacle and shows a pH optimum of 4.0 for both ssRNA and ssDNA (Yu et al. 2023), suggesting a protein adaptation to the acidic digestive fluids. DAN1 possesses all convergent amino acid substitutions found in a pair of CfENDO2 and ancestral ENDO2 sequences of Caryophyllales carnivorous plants, except for E167D (supplementary fig. S4). In addition to these convergent sites, CfENDO2 and NgENDO2 accumulated more convergent substitutions. These additional substitutions may refine the enzymatic activity of the ENDO2 protein further for the pitfall trap.

**Fig. 5.**
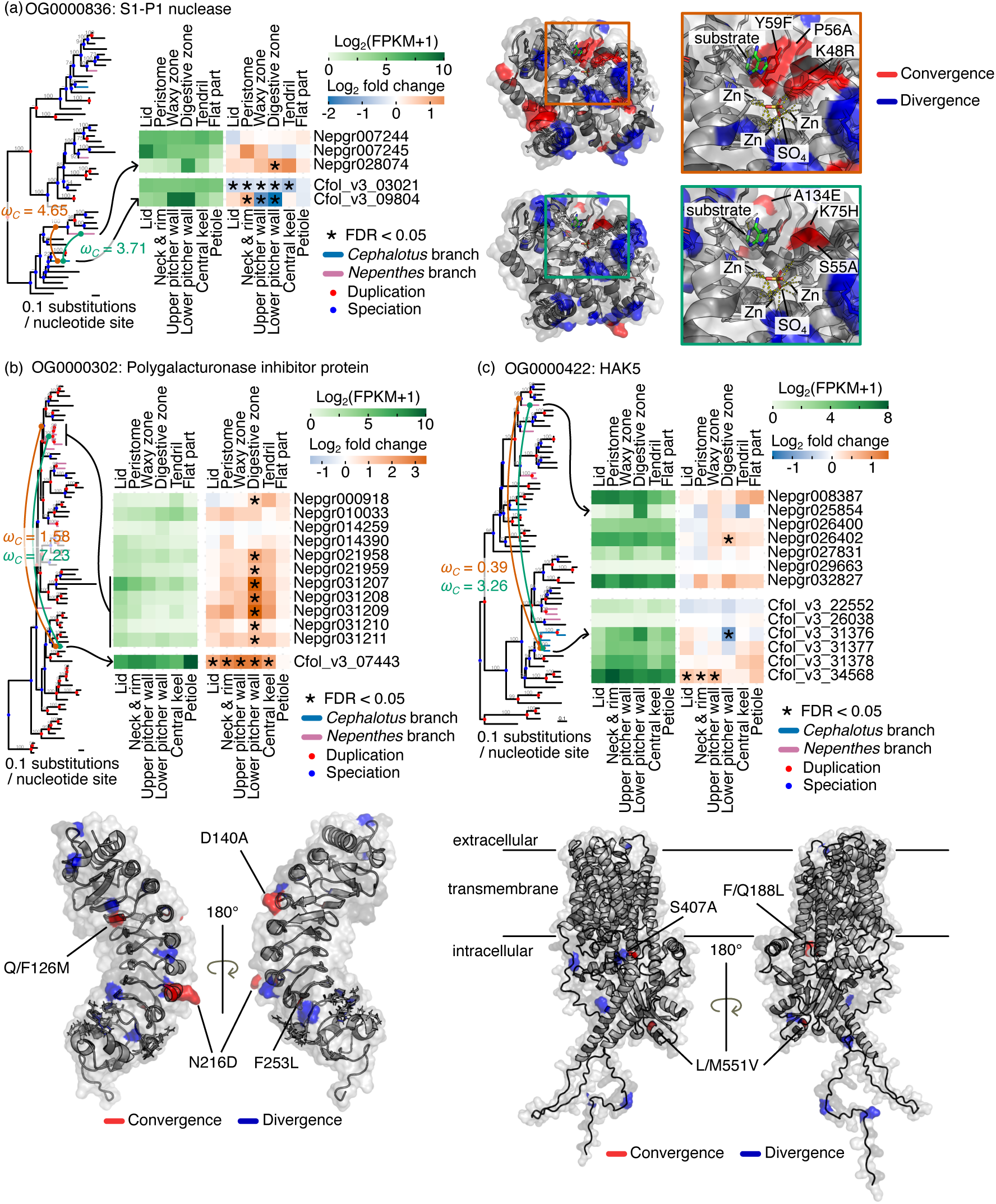
Convergent amino acid substitutions in proteins expressed in the gland-containing tissues. S1-P1 nuclease (a), Polygalacturonase-inhibiting protein (PGIP) (b), and K^+^ transporter HAK5 (c) are shown. *Cephalotus* and *Nepenthes* lineages are highlighted in blue and purple, respectively. Green lines connect focal branch pairs that exhibit high convergence rates, while brown lines link the focal *Cephalotus* branches to the stem branches of carnivorous Caryophyllales orthologs. The corresponding *ω_C_* values are indicated in matching colors. Ultrafast bootstrap values from IQ-TREE are indicated above the branches. Branches reconciled by GeneRax are marked with a hyphen (–). Node colors represent inferred evolutionary events: speciation (blue) and gene duplication (red). Expression levels (log₂(FPKM + 1)) under the basal condition and log₂ fold changes upon the feeding treatment are shown as heatmaps. Asterisks denote significant differential expression (FDR < 0.05). Convergent and divergent amino acid substitutions are mapped onto the protein structures and highlighted in red and blue, respectively. The protein structures were obtained from the Protein Data Bank entry or the AlphaFold Protein Structure Database (a, 3W52; b, 6W78; c, AF-Q5JK32-F1-v4). Site numbers correspond to positions in the protein structures. Note that the predicted HAK5 protein structure has very low pLDDT scores (< 50) in the cytoplasmic N-terminal (residues 1–55) and C-terminal regions (residues 670–706), indicating lower confidence in those areas. Black lines are drawn to indicate extracellular, transmembrane, and intracellular regions, but these are intended only as rough guidelines and do not reflect precise structural boundaries. For S1-P1 nuclease (a), two separate mapping patterns are displayed: one for the focal *Cephalotus*–*Nepenthes* branch pair, and another for the *Cephalotus* branch paired with the stem branch of carnivorous Caryophyllales ortholog. Magnified views of the substrate-binding sites are provided, with frame colors corresponding to the line colors linking branches in the tree. Substrate adenine is shown as green sticks. Only convergent substitutions near the substrate-binding site are labeled.

PGIPs have been detected in the digestive fluids of *Di. muscipula* and *Dr. adelae* (Schulze et al. 2012; Arai et al. 2021), suggesting a potential role in plant carnivory. In our analysis, we identified excess convergent amino acid substitutions in PGIPs, whose transcripts were upregulated following the feeding treatment in the two pitcher plant lineages (Fig. 5b). Notably, convergent substitutions were detected between the stem branch of five tandemly duplicated *N. gracilis PGIP* genes (Nepgr031207 to Nepgr031211) and a *C. follicularis PGIP* gene (Cfol_v3_07443), but not in the branch pair involving the common ancestor of carnivorous Caryophyllales. This pattern suggests a pitcher trap-specific adaptation rather than the initial establishment of carnivory. *Nepenthes PGIP* genes exhibited digestive zone-specific upregulation, whereas a *Cephalotus PGIP* gene was upregulated across all pitcher tissues except the petiole. Among the detected convergent substitutions, Q/F126M (corresponding to Y105 in *Phaseolus vulgaris* PGIP2) was located at the interaction site with fungal polygalacturonase (Xiao et al. 2024), potentially altering target specificity or binding affinity, or both. The remaining two convergent sites were positioned on the opposite side of the interaction interface, suggesting that they may influence other protein properties. Antifungal proteins are thought to prevent fungal growth in the digestive fluids, thereby reducing competition for nutrients (Buch et al. 2013). Thus, these amino acid substitutions, found exclusively in pitcher plants rather than across the entire carnivorous clade, may optimize PGIP function for the unique digestive environment of the pitfall trap.

HAK5, a potassium/proton symporter, has been previously reported in *Di. muscipula*, where its gene expression is induced by coronatine application and facilitates the absorption of prey-derived potassium into gland cells (Scherzer et al. 2015). In our analysis, we identified convergent amino acid substitutions between a *N. gracilis* HAK5 ortholog (Nepgr025854) specifically expressed in the digestive zone and a *C. follicularis* HAK5 ortholog (Cfol_v3_31376) specifically expressed in the lower pitcher wall, even though there are a total of five lineage-specific duplicates in the same clade (Fig. 5c). Nepgr025854 showed no significant response to the feeding treatment, whereas Cfol_v3_31376 was downregulated after the feeding treatment (Fig. 5c). Two detected convergent sites (F/Q188L and S407A) were in helices in the transmembrane part and one convergent site (L/M551V) was found in the cytosolic C-terminal domain. These substitution sites were not located at previously identified key residues essential for potassium binding in the transmembrane part (Tascón et al. 2020; Maierhofer et al. 2024) nor overlap with previously characterized auto-inhibitory or activation domains at the C-terminus (Ródenas et al. 2021), rendering them attractive candidate sites for future biochemical characterization.

Our analysis newly identified additional cases of convergent amino acid substitutions in independently evolved pitcher plants, expanding upon the previously recognized examples of digestive fluid proteins. Notably, the newly identified proteins include both secreted and non-secreted proteins, suggesting that selective pressures extend beyond extracellular proteins to also influence membrane-anchored and intracellular proteins that likely function within gland cells. Both *Nepenthes* gland cells and *Cephalotus* small glands possess permeable cuticles (Adlassnig et al. 2012), implying that gland cells are susceptible to external disturbances, which may have driven the adaptive evolution of non-secreted proteins responsible for maintaining cellular integrity, regulating ion transport, or protecting against damage from autodigestion.

## Conclusion

While both *C. follicularis* and *N. gracilis* upregulate protein synthesis in response to prey capture, their transcriptional responses are spatially distinct. In *C. follicularis*, DEGs are detected across all tissues, whereas in *N. gracilis*, DEGs are detected only in the digestive zone and the tendril (Saul et al. 2023). This suggests that pitcher tissues are more extensively differentiated in *Nepenthes* than in *Cephalotus*, exhibiting more compartmentalized responses in digestion-related tissues. Supporting this, a greater proportion of prey-derived nitrogen is found outside pitcher tissues in *N. mirabilis* (61.5%) compared to *C. follicularis* (26.1%) (Schulze et al. 1997). This conversely indicates that the *C. follicularis* pitchers exhibit less specialization for carnivory despite their extensive morphological adaptation. Higher photosynthetic activity of *C. follicularis* pitchers than *Nepenthes* pitchers corroborates this idea (Pavlovič 2011), yet upon prey capture, *C. follicularis* shifts the gene regulation toward carnivory by decreasing the expression of the photosynthetic-related genes (Fig. 1e). It is intriguing to investigate the tissue-wise feeding response in Sarraceniaceae because they rely more on prey-derived nitrogen (76.4% in *Darlingtonia*) but have a high photosynthetic activity comparable with *C. follicularis* in pitcher leaves (Pavlovič et al. 2007; Karagatzides and Ellison 2009).

The degree of specialization toward carnivory in the pitcher leaves of *N. gracilis* and *C. follicularis* is likely shaped by their distinct habitats, which differ markedly in climate and resource availability. *N. gracilis* thrives in warm, humid tropical regions where consistently high temperatures support abundant insect activity year-round (Anu et al. 2009; Schultheiss et al. 2022; Van Dijk et al. 2024). In such environments, strong selective pressures likely drive the evolution of highly specialized pitcher leaves. By contrast, *C. follicularis*, native to Albany, Australia, inhabits a region with greater seasonal temperature variation (Fukushima et al. 2021). This species displays developmental plasticity that allows it to produce pitcher or flat leaves in response to environmental cues, indicating a flexible reliance on carnivory. In its temperate habitat, where insect availability fluctuates depending on the time of the season, the benefits of strict specialization may not be substantial. Thus, habitat-driven differences in prey availability may have influenced the evolutionary trajectories of these two pitcher plant lineages, determining their varying degrees of specialization in carnivory.

## Materials and Methods

### Plant materials and growth conditions

Plants of *C. follicularis* were vegetatively propagated from the strain whose genome was sequenced previously (Fukushima et al. 2017), and grown in the half-strength MS medium supplemented with 3% sucrose, 1× Gamborg’s vitamins, 0.1% 2-(N-morpholino)ethanesulfonic acid, 0.05% Plant Preservative Mixture (Plant Cell Technology), and 0.3% Phytagel at 25 °C under long-day conditions (16 h light / 8 h dark).

### Tissue-specific RNA-seq with the mealworm extract treatment

One hundred mg of dried mealworms (batch number L400518, MultiFit Tiernahrungs GmbH) were powdered with mortars and pestles. The mealworm powder was dissolved in 1 ml of MilliQ water and vigorously vortexed. After the brief centrifugation, the supernatant was used for sample treatment. Approximately 10-30 μl of the mealworm extracts or Milli Q water (control) was supplied with open pitchers depending on the size of the pitcher. After 24 hours, pitchers were sampled and dissected as described in Fig. 1c using small blades, then snap-frozen in liquid nitrogen and stored at −80 °C. Dissecting one pitcher took approximately 90 seconds. The number of pitchers per replicate was 30-36. Total RNA was extracted using PureLink Plant RNA reagent (Life Technologies) according to the manufacturer’s instructions, and then further cleaned using the RNeasy Mini column (Qiagen). During cleanup, on-column DNA digestion was also performed using DNase I according to the manufacturer’s instructions (Qiagen). Library preparation and sequencing (paired-end, 150 bp) on NovaSeq 6000 (Illumina) were performed by Novogene.

### Differential gene expression analysis and gene ontology enrichment analysis

RNA-seq reads were quality-filtered using fastp v0.23.2 (Chen et al. 2018) with default parameters and pseudo-aligned to the *C. follicularis* transcriptome (Fukushima et al. 2017) using kallisto v0.48.0 (Bray et al. 2016) with default settings. Cross-species trimmed mean of M-values (TMM) normalization of the count was performed using AMALGKIT v0.12.3 (https://github.com/kfuku52/amalgkit) (Fukushima and Pollock 2020). The obtained FPKM values were used to plot heatmaps. DEGs were detected by comparing control and mealworm extract-treated samples for each tissue using the TCC package v1.36.0 (Sun et al. 2013), which internally utilizes edgeR v4.0.16 and TMM-normalization (Robinson et al. 2010). Low expression genes were excluded using the filterLowCountGenes function before TMM-normalization. Read count data for *N. gracilis* were obtained from the previous publication (Saul et al. 2023), and DEG analysis was performed using the same procedure as for *C. follicularis*. GO enrichment analysis was conducted with clusterProfiler v4.10.0 (Wu et al. 2021). GO terms for *C. follicularis* genes were assigned using EnTAP v0.10.8 (Hart et al. 2020), with homology searches performed against the TrEMBL, SwissProt (https://www.uniprot.org/), and NCBI plant RefSeq (https://ncbi.nlm.nih.gov/refseq/) databases. Annotations were selected based on Viridiplantae or Eukaryotes classifications in EggNOG.Tax.Scope. GO term information for *N. gracilis* was sourced from the previous publication (Saul et al. 2023).

### SOM clustering

SOM clustering was performed with the kohonen package v3.0.12 (Wehrens and Buydens 2007). Genes with the top 10% coefficient variation in TMM-normalized FPKM values were used for SOM clustering. FPKM values were mean-centered and variance-scaled, then used for SOM clustering in a 3 × 4 grid in hexagonal topology.

### Species tree inference

First, we inferred a phylogenetic tree of 15 species including *C. follicularis* and Caryophyllales carnivorous plants (Fig. 1a) (supplementary table S11) following the essentially same procedures used in the previous publication (Saul et al. 2023). Briefly, common single copy BUSCO genes (v5.3.2) (embryophyta_odb10) (Manni et al. 2021) among 15 species were aligned in-frame with mafft v7.508 (Katoh and Standley 2013) and tranalign from EMBOSS v6.6.0.0 (Rice et al. 2000). After trimming with trimAl v1.4.rev15 (Capella-Gutiérrez et al. 2009), all gene alignments were concatenated and used for maximum likelihood tree reconstruction under the GTR+R4 model with IQ-TREE2 v2.2.0.3 (Nguyen et al. 2015). The obtained tree was rooted using *Amborella trichopoda* as an outgroup and had the same topology as the previous publication (Saul et al. 2023). Next, divergence time estimation was performed with MCMCtree from the PAML package v4.9 (Yang 2007) using the same parameters as the previous publication (Saul et al. 2023). The reconstructed species tree and concatenated codon alignment were used as inputs. Branch lengths and substitution model parameters were initially estimated using BASEML with a global clock and the GTR+G model.

### Orthogroup classification

Orthogroup classification was performed with OrthoFinder v2.5.4 (Emms and Kelly 2019) using the inferred species tree as a guide. We selected orthogroups that included genes from at least 30% of the species in our dataset, resulting in a total of 13,023 orthogroups for downstream analyses. The hierarchical orthogroup classification at the node representing the most recent common ancestor of *C. follicularis* and *N. gracilis* was used to assess shared DEGs and SOM cluster members between the two species (Figs. 2,3 and supplementary figs. S1,3).

### Gene tree inference

For each orthogroup, a gene phylogenetic tree was inferred with IQ-TREE2 v2.2.0.3 under the GTR+G4 model using in-frame aligned and trimmed CDS sequences with mafft v7.508 (Katoh and Standley 2013), tranalign from EMBOSS v6.6.0.0 (Rice et al. 2000), and ClipKIT v1.3.0 (Steenwyk et al. 2020). Stop codons and ambiguous codons were replaced with gaps, and codon sites with many gaps were removed using ‘mask’ and ‘hammer’ functions in CDSKIT v0.10.2, respectively (https://github.com/kfuku52/cdskit). Obtained gene phylogenetic trees were reconciled with the inferred species tree using GeneRax v2.0.4 (--rec-model “UndatedDL”) (Morel et al. 2020). Divergence times of individual species-tree-aware gene phylogenetic trees were estimated with RADTE v0.2.0 (https://github.com/kfuku52/RADTE) using the dated species tree as a reference (Fukushima and Pollock 2020). Branching events in the gene trees were classified as either speciation or duplication using the species-overlap method (Huerta-Cepas et al. 2007).

### Jaccard similarity coefficient calculation

Jaccard similarity coefficient was calculated by dividing the size of the intersection by the size of the union of two hierarchical orthogroup sets. *P* values were calculated as the probabilities where 10,000 sets of simulated values exceed observed values. Simulations were performed by permuting hierarchical orthogroups used to calculate the original Jaccard similarity coefficient. To ensure consistency, contradictory hierarchical orthogroup assignments (e.g., assigning the same hierarchical orthogroup to both upregulated and downregulated categories within the same tissue) were avoided during the simulations. Multiple testing corrections were performed using the p.adjust function with the Benjamini-Hochberg method in R v4.3.2.

### Convergent amino acid substitution detection

We selected orthogroups that include both *N. gracilis* and *C. follicularis* genes upregulated in the digestive zone or the lower pitcher wall (FDR < 0.05, log_2_ fold change > 1), or specifically expressed in the gland-containing tissues (SOM cluster 4 in Fig. 3 for *N. gracilis* genes, and SOM cluster 1 and 5 in Fig. 3 for *C. follicularis* genes). In this analysis, OrthoFinder’s non-hierarchical orthogroups were used instead of hierarchical orthogroups to encompass distantly related branch pairs within the same orthogroup. By using non-hierarchical orthogroups, we detected excess convergence in key proteins such as HAK5, where the convergent branch pair split into two separate hierarchical orthogroups. CSUBST v1.1.4 estimated protein convergence for each selected orthogroup based on a codon alignment, a dated and reconciled gene phylogenetic tree, and a reconstructed ancestral codon state (Fukushima and Pollock 2023). Ancestral codon reconstructions were performed with IQ-TREE2 v2.2.0.3 using the “--ancestral” option (Nguyen et al. 2015).

### Homologous gene search

Homologous genes were searched by running TBLASTX v2.13.0 with an E-value threshold of 0.01 against 15 species CDS sequences used in the species tree inference. Gene phylogenetic trees were inferred through the same procedure used for the orthogroup analysis as described above.

## Data Availability

RNA-seq reads are available at the DNA Data Bank of Japan (DDBJ) under BioProject number PRJDB15743 (supplementary table S12). Other data are available in the article, its supplementary materials, or on Figshare (10.6084/m9.figshare.29410970).

## Code Availability

Scripts used in this study are available on Figshare (10.6084/m9.figshare.29410970).

## Acknowledgments

We acknowledge the following sources for funding: the Humboldt postdoctoral research fellowship (to T.W.), the Sofja Kovalevskaja programme of the Alexander von Humboldt Foundation (to K.F.), Deutsche Forschungsgemeinschaft research grants (454506241 to K.F.), a Human Frontier Science Program Young Investigators grant RGY0082/2021 (to K.F.), and JSPS KAKENHI 23K20050 (to K.F.). Computations were partially performed on the National Institute of Genetics supercomputer. This manuscript was written with the assistance of ChatGPT-4o.

## Author Contributions

T.W. and K.F. conceptualized the study. K.F. provided plant materials. T.W. conducted the experiments. T.W. analyzed the data. K.F. provided the in-house analysis pipeline. T.W. and K.F. wrote the manuscript.

## Supplementary Information for

## List of Supplementary Materials

supplementary texts 1-3

supplementary tables S1-12 (separate file)

supplementary figs. S1–6

supplementary references

supplementary dataset (separate file on Figshare: 10.6084/m9.figshare.29410970)

## supplementary texts

**supplementary text 1**. Potential role of immune elicitors in the constitutive expression of digestive fluid protein genes in *Cephalotus*. *Cephalotus* plants grown axenically, and therefore free of biotic stimuli, still show constitutive expression of digestive-fluid protein genes. Many of these genes are evolutionarily derived from defense-related loci (Fukushima et al. 2017), so the mechanism sustaining their expression remains unclear. One possible explanation is the continuous production of immune elicitors, such as *PATHOGENESIS-RELATED PROTEIN 1* (*PR1*), by apoplastic proteases including *CONSTITUTIVE DISEASE RESISTANCE* (*CDR1*). For example, in *Solanum lycopersicum*, CAPE1 peptide derived from a PR1 protein called SlPR1b induces the expression of defense response genes, including members of PR protein families (Chen et al. 2014). In *A. thaliana*, *XYLEM CYSTEINE PEPTIDASE 1* (*XCP1*) is recognized as a main protease responsible for producing AtCAPE9 peptide from PR1 proprotein (Chen et al. 2023). Notably, one *Cephalotus PR1* gene was preferentially expressed in the upper and lower pitcher walls, mirroring the expression pattern of genes encoding digestive fluid proteins (supplementary fig. S5). This gene harbors the “CNYD” substrate motif targeted by cysteine peptidase at its C-terminal. Although we did not observe clear tissue specificity, multiple cysteine peptidases were expressed highly across pitcher tissues (supplementary fig. S6). This suggests the constant production of the CAPE peptide from PR1 protein via protease activity, potentially eliciting a sustained immune response in the pitchers. Similarly, aspartic protease CDR1 is known to induce defense responses. In *A. thaliana*, *CDR1-D* mutant, which overexpresses *CDR1*, exhibits enhanced expression of *PR* genes (Xia et al. 2004). Its substrate is still unknown, but its functional conservation in rice has been reported (Prasad et al. 2009). Two *CDR1*-like genes in *C. follicularis* showed similar expression pattern to genes encoding digestive fluid proteins (supplementary fig. S7), suggesting a role in producing immune elicitors potentially linked to digestive fluid proteins.

**supplementary text 2**. **Feeding responses of jasmonic acid-related genes.** In the carnivorous Caryophyllales lineage, including *Nepenthes*, jasmonic acid (JA) signaling mediates feeding responses (Nakamura et al. 2013; Mithöfer et al. 2014; Buch et al. 2015; Böhm et al. 2016; Yilamujiang et al. 2016; Krausko et al. 2017). However, in *C. follicularis*, a member of the Oxalidales order, the endogenous JA level does not substantially change in response to prey capture, and coronatine, a bacterial toxin that mimics JA, does not induce digestive enzyme accumulation (Pavlovič et al. 2024). To further investigate the potential involvement of the JA signaling pathway in prey capture response in *C. follicularis*, we surveyed the transcriptional responses of genes involved in the JA biosynthesis and signaling (Wasternack and Song 2017; Wasternack and Feussner 2018) upon the feeding treatment. Two genes encoding allene oxide synthase (AOS), which convert lipoxygenase-derived hydroperoxides of linolenic and linoleic acids to precursors of JA, were upregulated after the feeding treatment and detected as DEGs in a few tissues (lid, neck, and upper pitcher wall), whereas there is no apparent induction of other JA biosynthesis genes (supplementary fig. S8a,b). In addition, *CORONATINE-INSENSITIVE 1* (*COI1*) and *JASMONATE-ZIM DOMAIN* (*JAZ*), encoding co-receptor complex of JA (Chini et al. 2007; Thines et al. 2007; Fonseca et al. 2009), were also not significantly induced except for one *JAZ* gene (Cfol_v3_19722), which showed only weak induction. In *N. gracilis*, three distinct genes encoding 12-oxophytodienoic acid (OPDA) reductase 3 (OPR3), required for JA biosynthesis, were specifically upregulated in the digestive zone. In contrast, other JA biosynthesis genes were not detected as DEGs across any tissues except for one gene encoding 13-lipoxygenase (LOX) which was instead downregulated (supplementary fig. S8). Signaling components were slightly upregulated in the lid and the waxy zone, but those changes were not significant.

Although no clear pattern emerged, our data indicate that JA-related genes are differentially regulated in both *Nepenthes* and *Cephalotus*, with over half of the analyzed *Cephalotus* genes showing significant regulation. Considering the stable abundance of digestive enzymes (Pavlovič et al. 2024), these findings suggest that JA contributes to feeding responses in *Cephalotus* beyond the secretion of digestive-fluid proteins. While carnivorous plants from different lineages employ a similar set of digestive enzymes repurposed from defense response (Fukushima et al. 2017), the transcriptional regulation of these enzymes varies across lineages as shown above and in previous research (Kocáb et al. 2020; Pavlovič et al. 2024), reflecting divergent evolutionary paths.

**supplementary text 3**. **Upregulation of protein synthesis may reflect nitrogen assimilation.** Upregulation of genes involved in protein synthesis is a major feeding response shared between the two pitcher plant lineages (supplementary table S5). In *N. gracilis*, this response is thought to be associated with the rapid production of digestive enzymes triggered by the feeding treatment (Saul et al. 2023). However, this scenario is not likely in *C. follicularis* because its digestive fluid proteins were rather downregulated after the feeding treatment (Fig. 4c). Alternatively, we propose that the upregulation of protein synthesis in *C. follicularis* may be linked to the process of nitrogen assimilation. In *A. thaliana*, nitrogen assimilation- and protein synthesis-related genes are quickly upregulated under nitrogen-replete conditions (Scheible et al. 2004). Similarly in *C. follicularis*, genes associated with the glutamine synthetase-glutamate synthase pathway, which is the first step of nitrogen assimilation (Masclaux-Daubresse et al. 2010), were upregulated particularly in the lower pitcher wall (supplementary fig. S9), potentially driven by prey-derived proteins or their degradation products such as ammonium.

## supplementary tables

**supplementary table S1.** Results of differential gene expression analysis across pitcher tissues in *C. follicularis*. The merged output of DEG analyses for all sampled tissues. Tissue identity is indicated by the suffix in column names, using the following abbreviations: LID, lid; NEC, neck and rim; UPP, upper pitcher wall; LWP, lower pitcher wall; KEL, central keel; PET, petiole.

- gene_id: Gene identifier.
- m.value: Log2 fold change between the two conditions.
- q.value: Adjusted p-value after multiple testing correction using the Benjamini-Hochberg false discovery rate.
- estimatedDEG: Binary indicator of differential expression status (1 = DEG, 0 = non-DEG).

**supplementary table S2.** GO enrichment analysis results of DEGs in each set association in the upset plot. Only GO terms with more than three associated genes and an adjusted q-value < 0.05 (Benjamini-Hochberg method) are included. Sheet names indicate gene set associations corresponding to those shown in Fig. 1. Tissue name abbreviations follow the same format as described in supplementary table S1.

**supplementary table S3.** Results of differential gene expression analysis across pitcher tissues in *N. gracilis*. The merged output of DEG analyses for all sampled tissues. Tissue identity is indicated by the suffix in column names, using the following abbreviations: LID, lid; PRS, peristome; WXZ, waxy zone; DGZ, digestive zone; TDR, tendril; FLP, flat part.

**supplementary table S4**. Orthologous gene group classification by OrthoFinder. The column labeled “HOG” indicates the hierarchical orthogroup (HOG) assignments determined at the node representing the most recent common ancestor of *C. follicularis* and *N. gracilis*.

**supplementary table S5.** GO enrichment analysis results of DEGs in commonly upregulated or downregulated orthogroups in the lower pitcher wall of *C. follicularis* pitcher and the digestive zone of *N. gracilis* pitcher. Only GO terms with more than three associated genes and an adjusted q-value < 0.05 (Benjamini-Hochberg method) are included. Sheet name includes abbreviations as follows: Cf, *Cephalotus follicularis*; Ng, *Nepenthes gracilis*. Tissue name abbreviations follow the same format as described in supplementary table S1 and supplementary table S3.

**supplementary table S6.** SOM clustering results based on the gene expression dataset before the feeding treatment. Each gene is assigned to a cluster, indicated in the “unit.class” column. Mean expression values for each tissue are provided. “Zmean” columns represent Z-scores calculated from the mean expression values across tissues. Tissue names follow the abbreviations used in supplementary table S1 and supplementary table S3. Treatments are abbreviated as follows: CON, control; WOR, mealworm extract.

**supplementary table S7.** GO enrichment analysis results of the SOM clusters in supplementary table S6. Only GO terms with more than three associated genes and an adjusted q-value < 0.05 (Benjamini-Hochberg method) are included. Sheet name includes abbreviations as follows: Cf, *Cephalotus follicularis*; Ng, *Nepenthes gracilis*.

**supplementary table S8**. SOM clustering results based on the gene expression dataset following the feeding treatment. Each gene is assigned to a cluster, indicated in the “unit.class” column. Note that the cluster numbers are independent of those in supplementary table S6. Mean expression values for each tissue are provided. “Zmean” columns represent Z-scores calculated from the mean expression values across tissues. Tissue names follow the abbreviations used in supplementary table S1 and Supplementary table S3. Treatments are abbreviated as follows: CON, control; WOR, mealworm extract.

**supplementary table S9.** GO enrichment analysis results of the SOM clusters in supplementary table S8. Only GO terms with more than three associated genes and an adjusted q-value < 0.05 (Benjamini-Hochberg method) are included. Sheet name includes abbreviations as follows: Cf, *Cephalotus follicularis*; Ng, *Nepenthes gracilis*.

**supplementary table S10**. CSUBST analysis results with *ω_C_* > 3 and *O_C_^N^* > 3. Each row represents a result of one branch pair. The numerical suffix (e.g., _1 or _2) indicates whether the column pertains to branch_id_1 or branch_id_2.

- branch_id_1, branch_id_2: Identifiers for the focal branches being analyzed.
- omegaCany2spe: The rate of combinatorial codon substitutions from any ancestral codons to specific descendant codons (equivalent to *ω_C_* in the manuscript).
- OCNany2spe: The number of combinatorial substitutions (corresponds to *O_C_^N^* in the manuscript).
- spnode_coverage: species coverage on the branch
- sprot_recname: functional annotation from RPS-BLAST.

**supplementary table S11**. Genome information sources for the 15 species used in this study.

**supplementary table S12**. DDBJ entry information.

## supplementary figures

**supplementary fig. S1.**
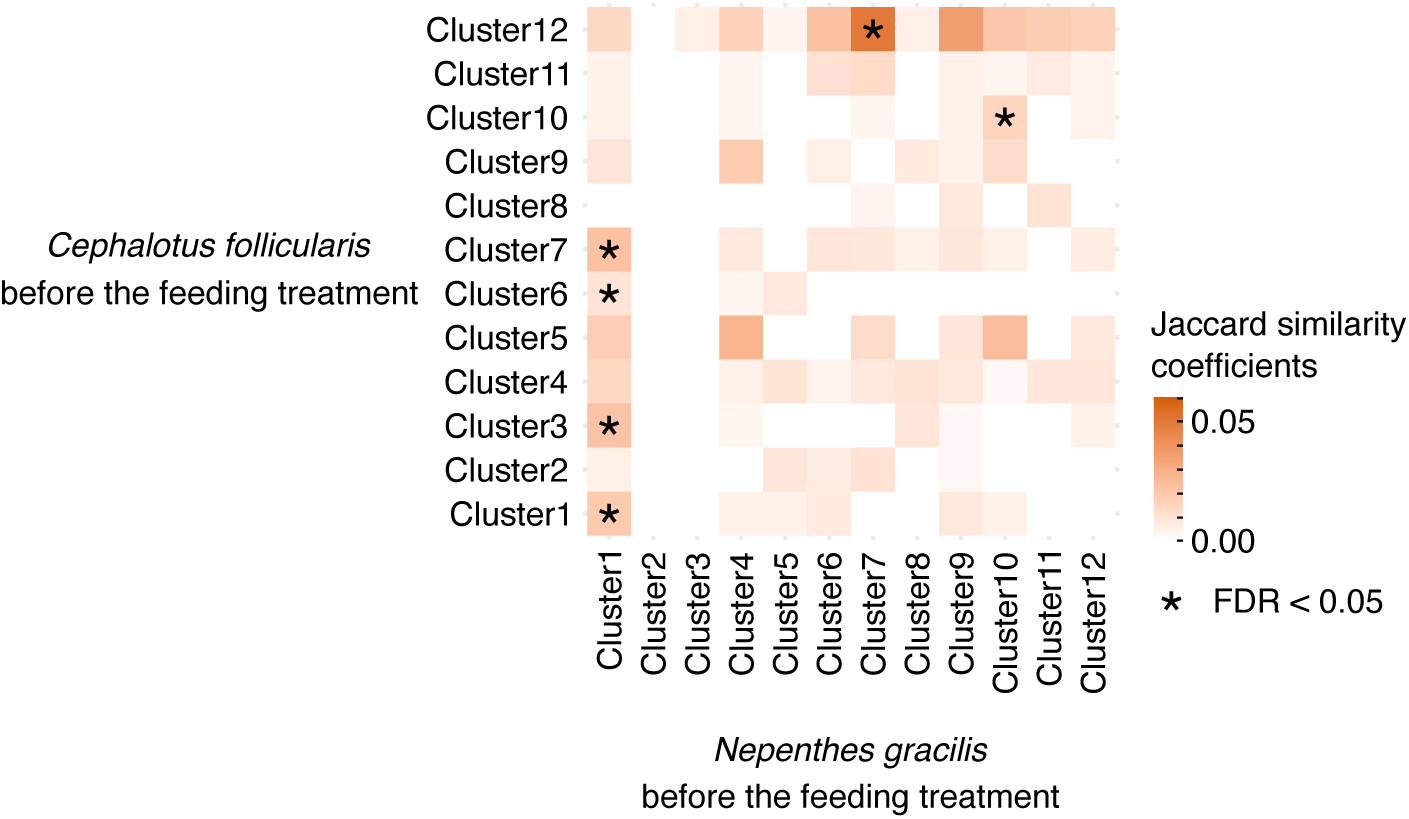
The Jaccard similarity coefficients (JC) between orthogroup sets from pairs of SOM clusters related to Fig. 3. Asterisks denote significantly high JC values (FDR < 0.05). *P* values were computed using permutation tests, and false discovery rates (FDRs) were adjusted using the Benjamini–Hochberg method for multiple testing correction.

**supplementary fig. S2.**
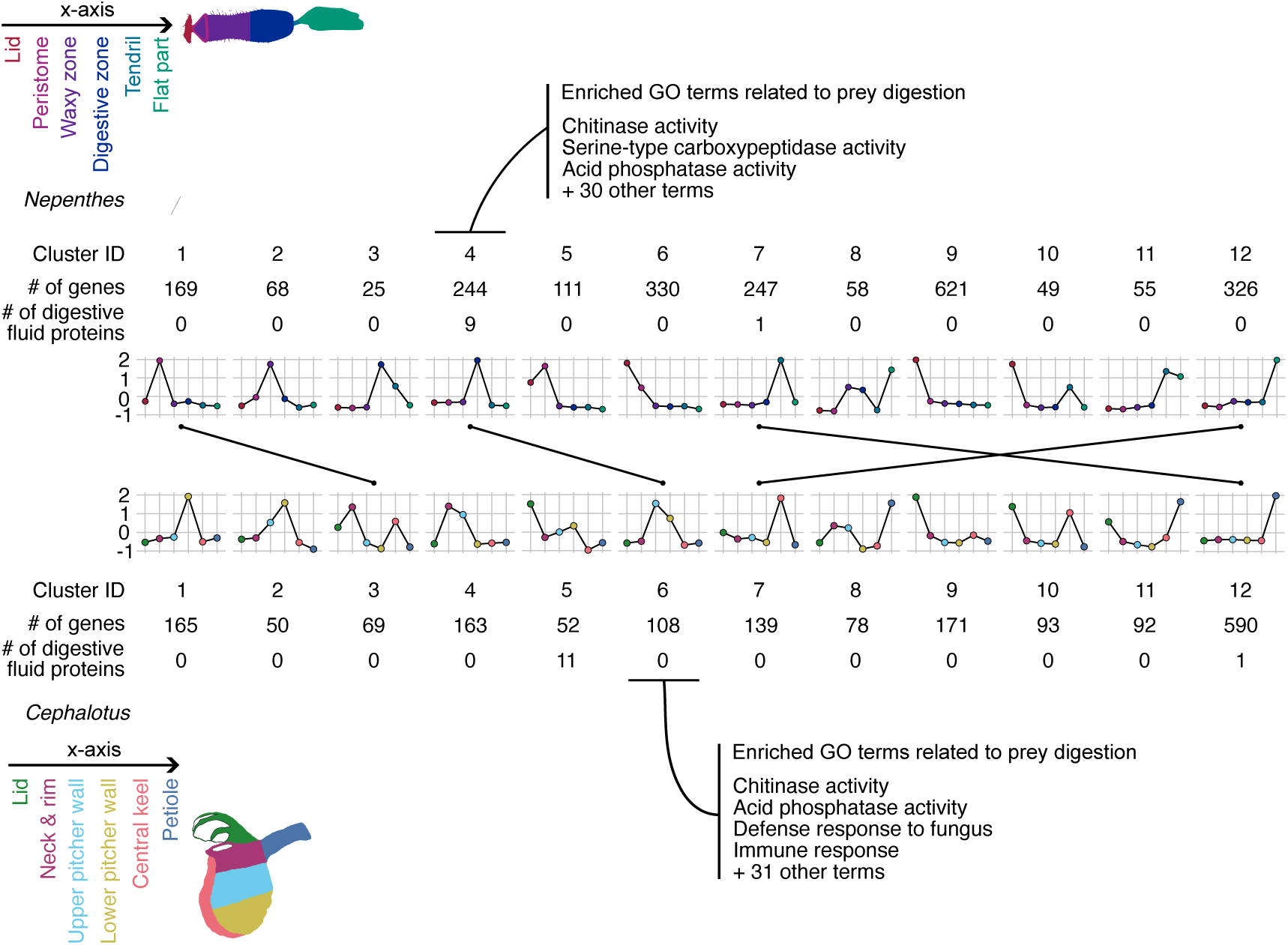
Comparison of tissue-specific gene expression following the feeding treatment in *N. gracilis* and *C. follicularis* pitchers. SOM clustering was performed on genes with high coefficient of variation in expression after the feeding treatment across six pitcher tissues in *N. gracilis* (top) and *C. follicularis* (bottom). The x-axis labels are shown at the top-left and bottom-left corners of the panels. The y-axis represents the average z-score of gene expression within each cluster. Clusters with significantly similar orthogroup compositions between the two species (Jaccard similarity coefficient, FDR < 0.05) are connected by black lines. Selected enriched GO terms for representative clusters are also shown. FDR values were calculated using the Benjamini–Hochberg method for multiple testing correction after *P* values were obtained through permutation tests.

**supplementary fig. S3.**
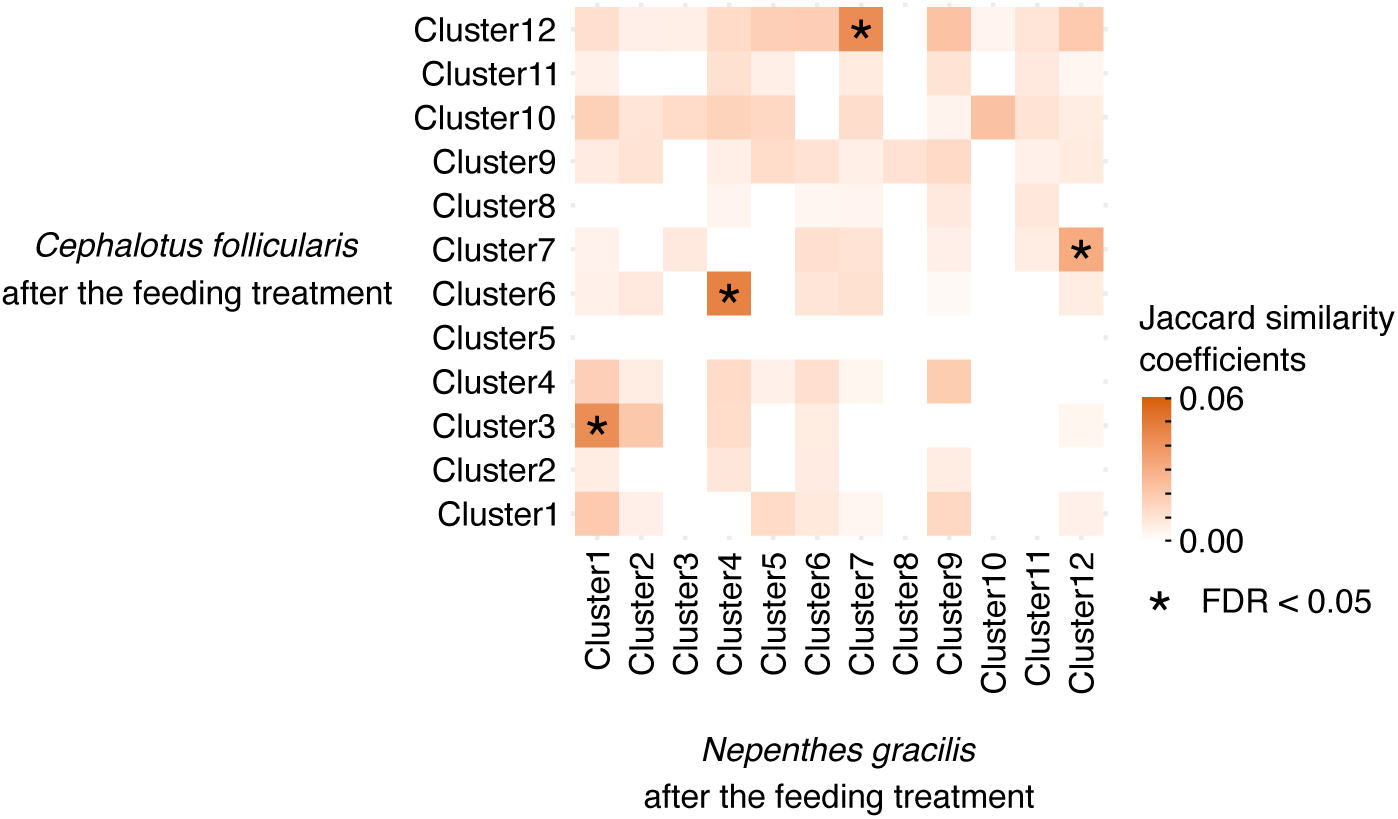
The Jaccard similarity coefficients (JC) between orthogroup sets from pairs of SOM clusters related to supplementary fig. S2. Asterisks denote significantly high JC values (FDR < 0.05). *P* values were computed using permutation tests, and false discovery rates (FDRs) were adjusted using the Benjamini–Hochberg method for multiple testing correction.

**supplementary fig. S4.**
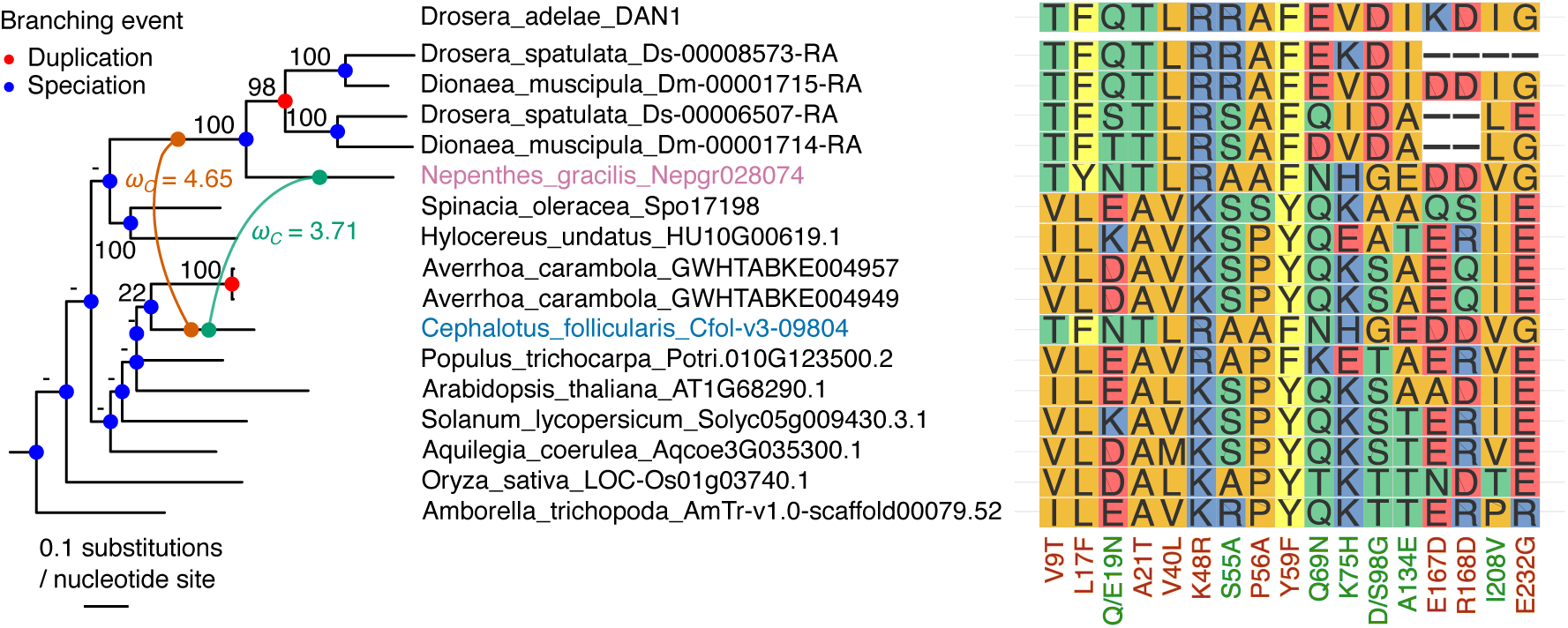
Convergent sites in ENDO2 protein, including *Drosera adelae* DAN1. *Cephalotus* and *Nepenthes* genes are highlighted in blue and purple, respectively. A green line connects the focal branch pair that exhibits a high convergence rate, while a brown line links the focal *Cephalotus* branch to the stem branch of carnivorous Caryophyllales ortholog. The corresponding *ω_C_* values are indicated in matching colors. Ultrafast bootstrap values from IQ-TREE are indicated above the branches. Branches reconciled by GeneRax are marked with a hyphen (–). Node colors represent inferred evolutionary events: speciation (blue) and gene duplication (red). Amino acids are depicted with colors corresponding to their side-chain chemistry. The x-axis label indicates substitution sites. The site numbers align with those found in the PDB entry (3W52), and font colors correspond to the branch pair in the phylogenetic tree.

**supplementary fig. S5.**
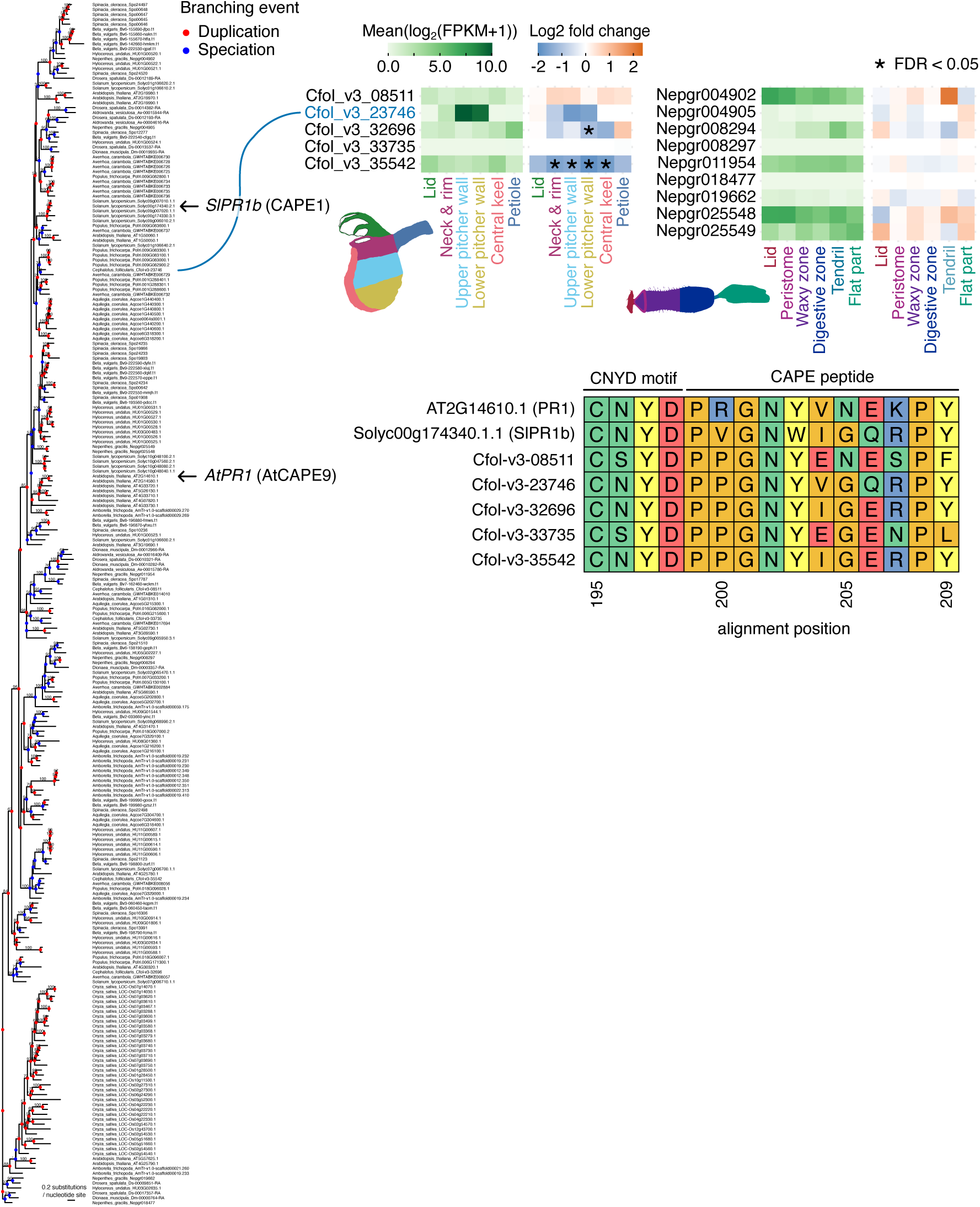
A *PR1*-like gene is specifically expressed in the upper and lower pitcher walls of *C. follicularis*. The phylogenetic tree highlights the positions of previously described *PR1* genes known to be precursors of CAPE peptides (indicated by arrows). Ultrafast bootstrap values from IQ-TREE are indicated above the branches. Branches reconciled by GeneRax are marked with a hyphen (–). Node colors represent inferred evolutionary events: speciation (blue) and gene duplication (red). Heatmaps display FPKM values under control conditions and fold changes following feeding treatment in the pitchers of *C. follicularis* and *N. gracilis*. A gene exhibiting expression patterns similar to digestive fluid proteins (corresponding to SOM cluster 5 in Fig. 3) is marked in blue. Asterisks denote significant differential expression (FDR < 0.05). The amino acid alignment illustrates the conservation of the “CNYD” motif and CAPE peptide sequences at the C-terminus of PR1 proteins. Amino acids are color-coded according to their side-chain chemical properties.

**supplementary fig. S6.**
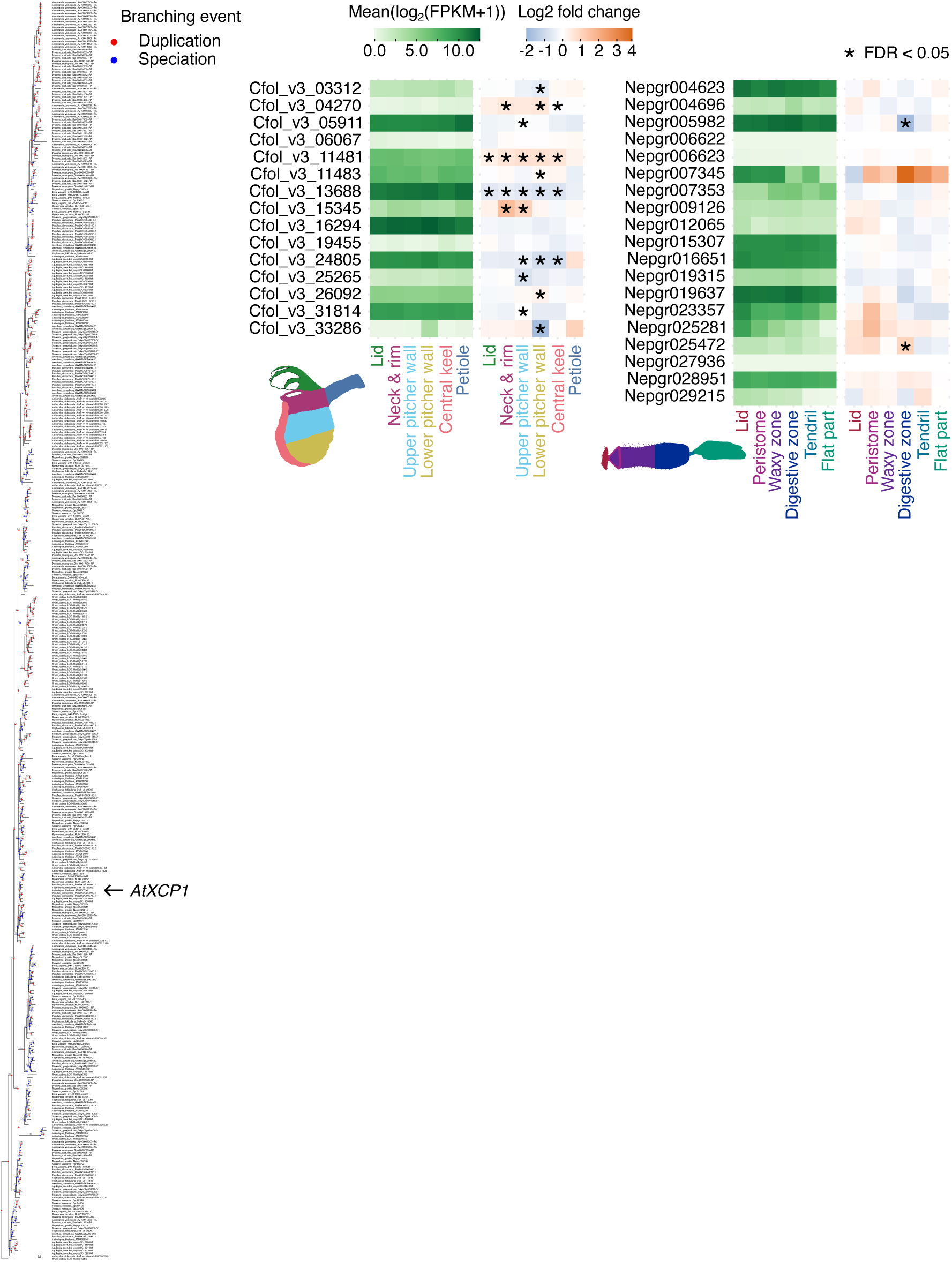
Papain-like cysteine proteases expression in the *C. follicularis* and *N. gracilis* pitcher. The phylogenetic tree highlights the position of *AtXCP1* known to produce CAPE peptides (indicated by an arrow). Ultrafast bootstrap values from IQ-TREE are indicated above the branches. Branches reconciled by GeneRax are marked with a hyphen (–). Node colors represent inferred evolutionary events: speciation (blue) and gene duplication (red). Heatmaps display FPKM values under control conditions and fold changes following feeding treatment in the pitchers of *C. follicularis* and *N. gracilis*. Asterisks denote significant differential expression (FDR < 0.05).

**supplementary fig. S7.**
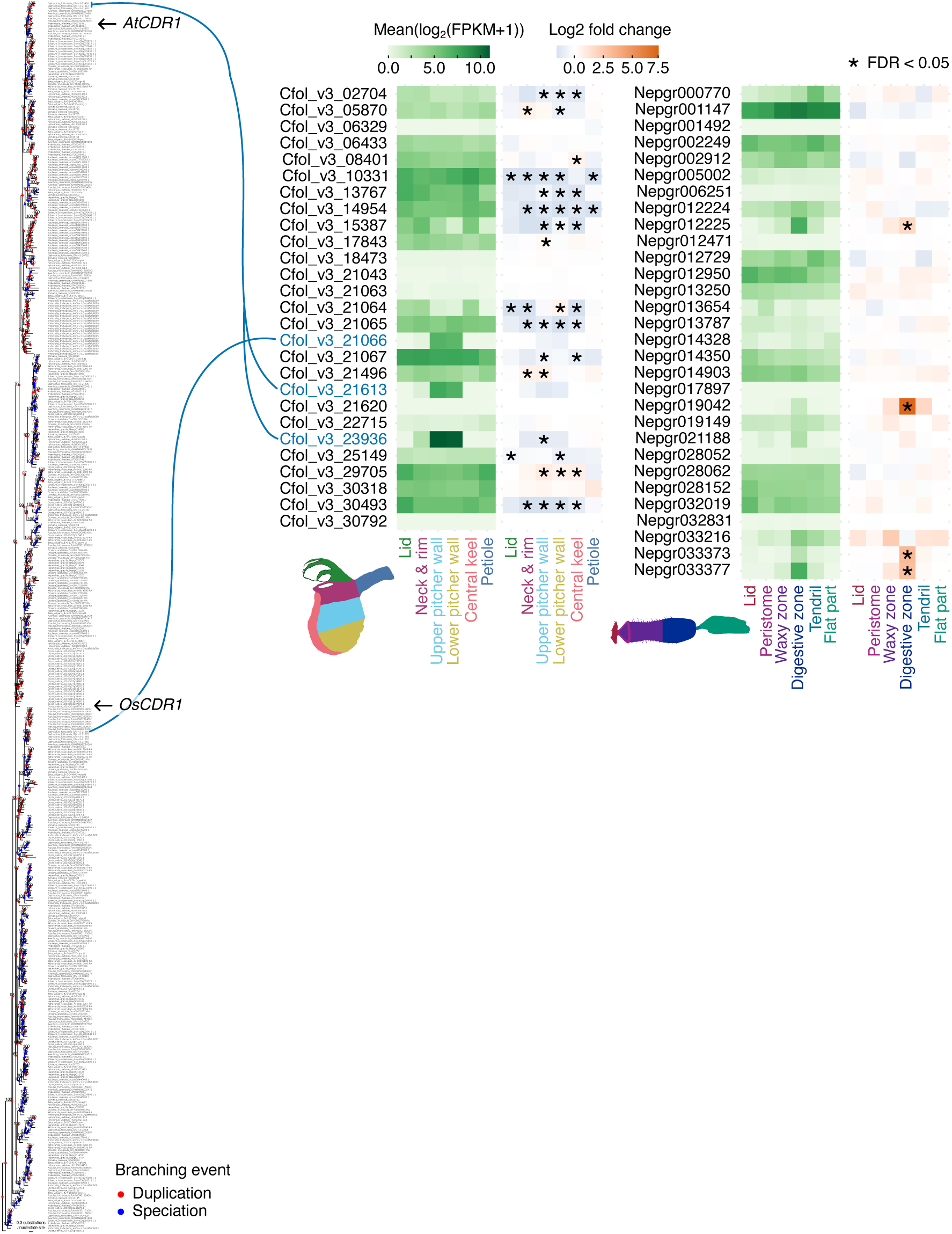
Expression profiles of aspartic proteases in the *C. follicularis* and *N. gracilis* pitcher. The phylogenetic tree highlights the positions of *CDR1* genes known to enhance immune responses (indicated by arrows). Ultrafast bootstrap values from IQ-TREE are indicated above the branches. Branches reconciled by GeneRax are marked with a hyphen (–). Node colors represent inferred evolutionary events: speciation (blue) and gene duplication (red). Heatmaps show FPKM values under control conditions and fold changes after feeding treatment in the pitchers of *C. follicularis* and *N. gracilis*. Genes exhibiting expression patterns similar to digestive fluid proteins (corresponding to SOM cluster 5 in Fig. 3) are marked in blue. Asterisks denote significant differential expression (FDR < 0.05).

**supplementary fig. S8.**
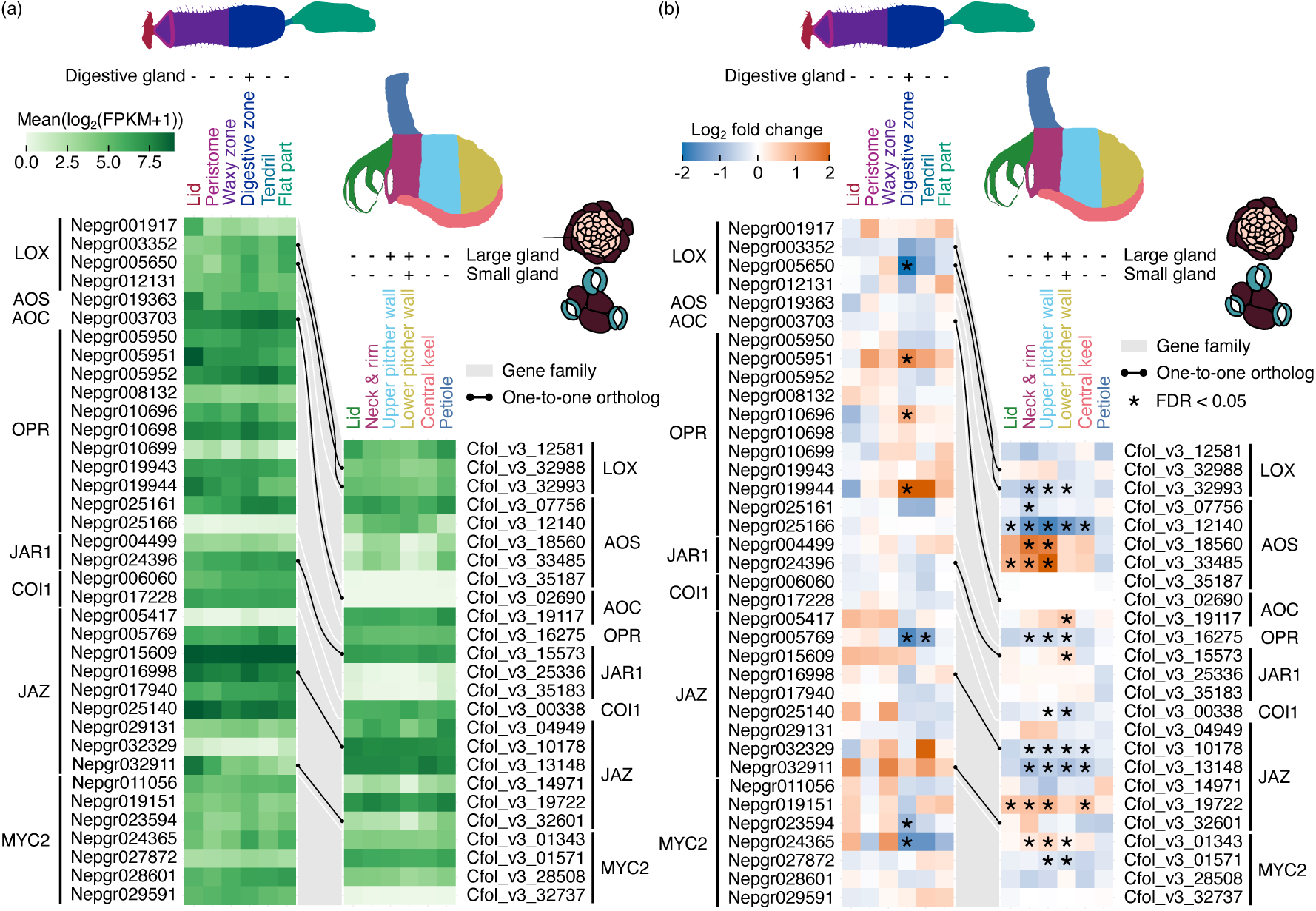
Transcriptional responses of jasmonic acid (JA)-related genes to the feeding treatment in *C. follicularis* and *N. gracilis* pitchers. (a) Expression level (log_2_(FPKM+1)) of the JA-related genes. (b) Log_2_-fold change of the JA-related genes upon the feeding treatment. Grey ribbons connect genes in the same gene family. The black dots and lines connect the orthologous genes. Asterisks in (b) indicate FDR < 0.05 in differential gene expression analysis.

**supplementary fig. S9.**
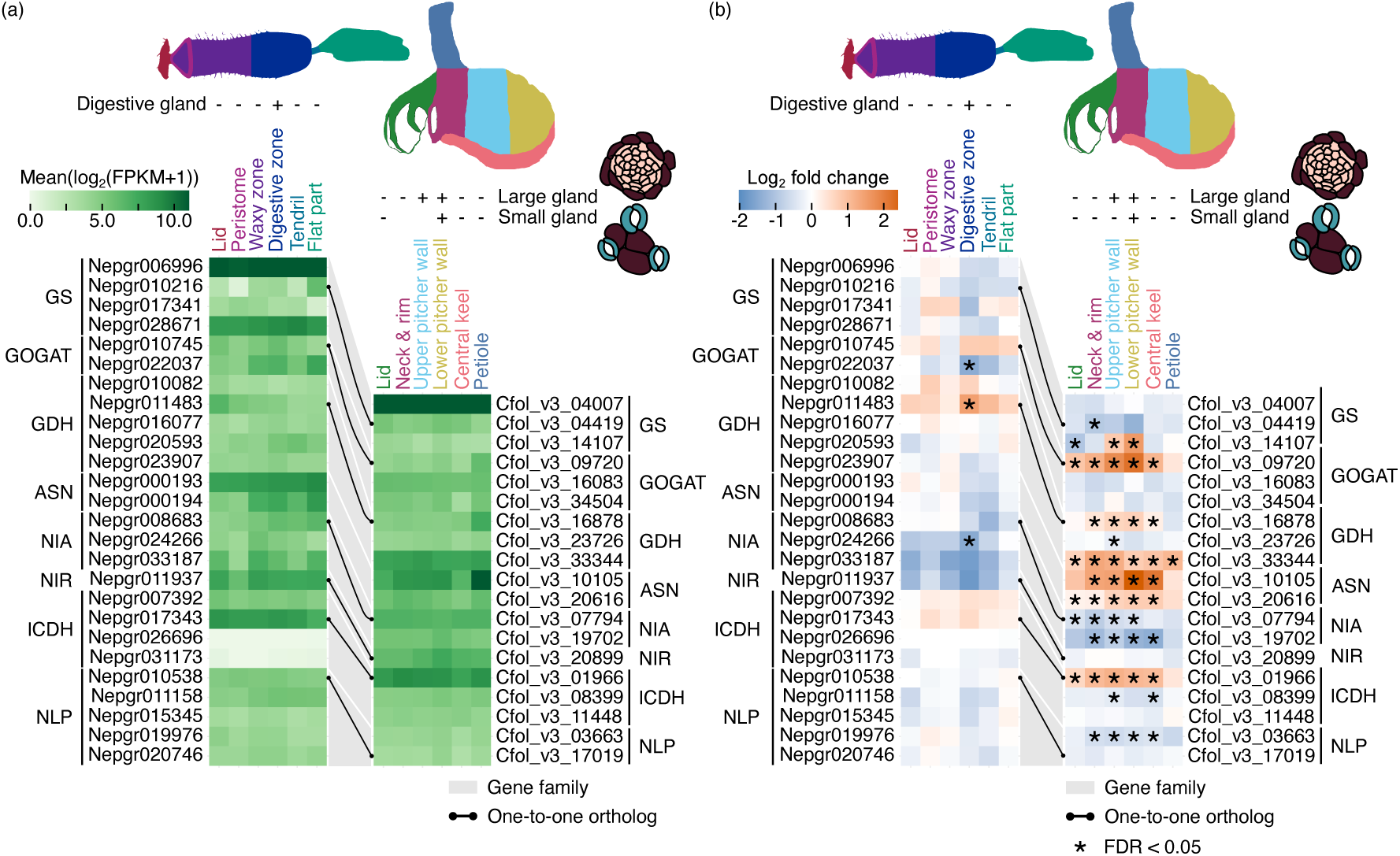
Transcriptional responses of nitrogen assimilation genes to the feeding treatment in *C. follicularis* and *N. gracilis* pitchers. (a) Expression level (log_2_(FPKM+1)) of the nitrogen assimilation genes. (b) Log_2_-fold change of the nitrogen assimilation genes upon the feeding treatment. Grey ribbons connect genes in the same gene family. The black dots and lines connect the orthologous genes. Asterisks in (b) indicate FDR < 0.05 in differential gene expression analysis.

